# Hierarchical Heterogeneities in Spatiotemporal Dynamics of the Cytoplasm

**DOI:** 10.1101/2025.05.08.652886

**Authors:** Conrad Möckel, Abin Biswas, Christine Schweitzer, Simone Reber, Aleksei Chechkin, Vasily Zaburdaev, Jochen Guck

## Abstract

The physical properties of the cytoplasm fundamentally constrain the dynamics and fidelity of cellular processes. Endogenous fluctuations carry signatures of these properties, yet how their collective dynamics across mesoscopic spatiotemporal scales relate to cytoplasmic heterogeneity and organization remains poorly understood. Here, we use differential dynamic microscopy (DDM) to quantify these fluctuations in eukaryotic cytoplasms. In *Xenopus laevis* egg extracts, we identify signatures of subdiffusive fractional Brownian motion (fBm) and non-Gaussian displacement distributions, consistent with a crowded, heterogeneous environment. We introduce an empirical model combining fBm with an inverse Gaussian diffusivity distribution, enabling robust estimation of fractional diffusivities and scaling exponents across conditions. Denser cytoplasm exhibits lower fractional diffusivity and scaling exponent, while energy addition induces non-stationary dynamics. Stabilization of microtubule networks introduces a secondary timescale, rationalized by a two-state fBm model separating cytosolic and network-associated contributions. In HeLa cells, the framework reveals comparable scaling exponents but lower diffusivities than in egg extract. Upon microtubule depolymerization, the dynamics become slower yet more Brownian-like. This highlights dual roles of the cytoskeleton as confinement but also as fluidizer of the cytoplasm. These results establish label-free DDM as an interpretable, collective, multiscale readout of how composition, activity, and cytoskeletal organization shape hierarchical cytoplasmic dynamics.

## I. INTRODUCTION

The cytoplasm is a highly dynamic and heterogeneous environment, comprising a complex mixture of macro-molecules, organelles, cytoskeletal networks, and phase-separated compartments [1–3]. The behavior of these components is shaped by physical properties such as local crowdedness, viscosity, and density, which collectively influence diffusion modes, affect interactions between specific macromolecular species, and ultimately drive biological functions [3–8]. However, the precise mechanisms governing cytoplasmic dynamics, such as anomalous diffusion [9], are not yet fully understood. Relating these dynamics to the underlying heterogeneity of the cytoplasm is therefore important for elucidating a wide range of biological phenomena, including self-organization, signaling, cell division, migration, homeostasis, disease, and morphogenesis [2, 10–13].

Experimentally characterizing cytoplasmic dynamics across length and time scales remains challenging [14]. Standard techniques, such as atomic force microscopy (AFM) [15], active and passive microrheology via single-particle tracking [16–20], fluorescence correlation spectroscopy (FCS) [21–23], and micropipette aspiration [24, 25], provide invaluable insights. However, these methods are often limited by localized measurements, the introduction of extrinsic labels or tracer particles, or reliance on theoretical constitutive models to infer subcellular properties from global cellular perturbations [17, 21, 26, 27].

In recent years, differential dynamic microscopy (DDM) has emerged as a complementary technique for inferring dynamical properties from microscopy images acquired with a contrast modality of choice [28, 29] (for a review, see [30]). When endogenous fluctuations generate measurable image contrast, DDM can provide a collective dynamical readout without the need for dedicated labels or tracer particles. Such label-free applications of DDM have been demonstrated in several biological systems, including active transport and advection in *Drosophila* ooplasm [31], microorganism motility [32, 33], protein size determination [34], and intestinal mucus dynamics [35]. In DDM, these collective dynamics are encoded in the intermediate scattering function (ISF), specifically its real part ℜ [*f* (*q*, Δ*t*)] [28, 29].

The ISF is the spatial Fourier transform of the van Hove correlation function *G*(Δ*r*, Δ*t*) and describes the temporal evolution of spatial correlations within the scattering population [36]. Experimentally, the ISF is inferred from differences in a series of digital microscope images *I*(***x***, *t*) in the spatial Fourier domain

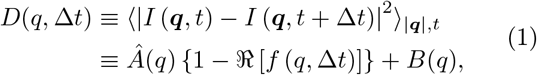

where *D*(*q*, Δ*t*) is the DDM observable and ⟨…⟩_|***q***|,*t*_ denotes the time and azimuthal average at the displacement mode *q* = |***q***| = 2*π/*Δ*r* (see Figure 1B). The amplitude term *Â*(*q*) ≡ 2⟨|*I*(***q***, *t*)|^2^⟩_|***q***|,*t*_−*B*(*q*) is related to the static structure factor *S*(*q*) as *Â*(*q*) ∝ | *T*(*q*) ^2^*S*(*q*), where *T*(*q*) is a setup-specific optical transfer function, and *B*(*q*) accounts for the noise characteristics of the camera [29, 37] (details in Figure 1 and *Supplementary Material*)^1^. The experimentally accessible time and length scales, set by the camera frame rate, optical magnification, and contrast generation method, determine which processes contribute to ℜ [*f* (*q*, Δ*t*)] and thus which dynamical information can be extracted.

**FIG. 1:**
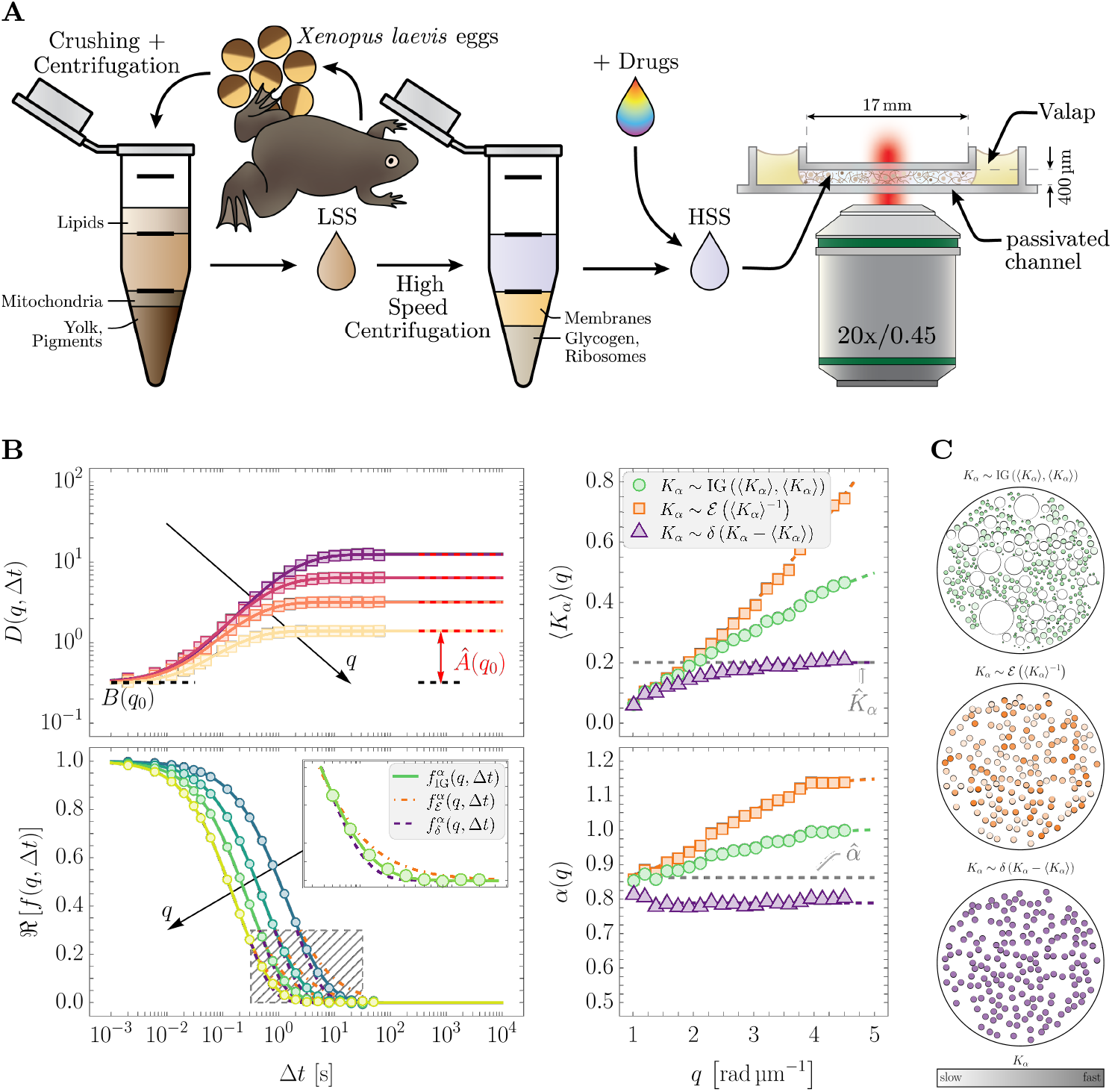
Schematic overview of the experimental setup and results. **A**: Production of high-speed *Xenopus laevis* egg extract (HSS) from low-speed (LSS) by high-speed centrifugation and loading into a microfluidic channel (dimensions not to scale). **B**: Left: Azimuthally averaged DDM map *D*(*q*, Δ*t*) over lag time Δ*t* (top), with black arrow showing dispersion over displacement modes *q*, black dashed line for camera noise *B*(*q*_0_), and red double arrow for amplitude estimate *Â*(*q*_0_) (see *Materials and Methods*) for *q*_0_ ≈ 4.13 rad µm^−1^; corresponding real part of the intermediate scattering function ℜ [*f* (*q*, Δ*t*)] (bottom) computed using Equation (1). Inset: Best fits of three closed-form models Equations (4) to (6) to data for *q*_0_ ≈ 3.44 rad µm^−1^ (see *Supplementary Material* for derivations). Right: Extracted best-fit parameters from the three models as a function of *q*; top, dimensionally normalized mean fractional diffusivity ⟨*K*_*α*_⟩(*q*); bottom, fractional exponent *α*(*q*). Colorful dashed lines are sigmoidal fits; gray dashed lines are inferred parameter estimates 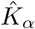 and 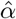 from Monte Carlo simulations (see *Supplementary Material*; Figure S3). **C**: Schematic representation of the model assumptions in **B**: top, scatterers with a size distribution and a distribution of (fractional) diffusivities following the generalized Stokes–Einstein relation described in the main text with constant fractional viscosity *η*_*α*_; middle, uniform size and exponential distribution of (fractional) diffusivities; bottom, uniform size and (fractional) diffusivity.

In this study, we use wide-field DDM to investigate the collective dynamical properties of the cytoplasm in a label-free manner. We first consider *Xenopus laevis* egg extract as a simplified model system for eukaryotic cytoplasm (Figure 1A), using bright-field microscopy. Egg extract is a highly dynamic and heterogeneous system, predominantly consisting of a complex mixture of proteins at physiological concentrations [22, 38]. The measured ISF of the extract exhibits characteristics of subdiffusive fractional Brownian motion (fBm) and non-Gaussianity, phenomena frequently associated with intracellular environments [23, 39–41]. We propose an empirical model that combines fBm with an inverse Gaussian distribution of diffusivities, motivated by the broad distribution of molecular sizes within the cytoplasm [38], and provides robust fits to our experimental data.

Our analysis captures shifts in cytoplasmic dynamics across a broad range of perturbations. Changing the composition toward a more concentrated extract reduces the scaling exponent of transport as well as the effective diffusivity. By enabling ATP-driven processes in the extract, we observe non-stationary dynamics with characteristic times of ~30 min. Furthermore, by stabilizing microtubules in the extract, we identify a second characteristic timescale in the ISF, revealing an additional hierarchy in the measured dynamics. We describe this behavior with a two-state fBm model that accounts for size-selective interactions between cytoplasmic scatterers and the microtubule network, allowing us to infer a characteristic crossover length scale comparable to the independently measured microtubule mesh size.

Finally, to connect these egg-extract-based findings to a cellular context, we show that the same DDM framework is readily applicable to quantify the properties of single HeLa cells using phase-contrast microscopy. Interestingly, microtubule perturbation with Nocodazole shifts the dynamics toward more Brownian-like transport scaling and substantially reduces the effective diffusivity. This suggests that microtubules affect the scaling and magnitude of intracellular fluctuations in distinct ways, consistent with a dual role in promoting subdiffusive dynamics while enhancing intracellular agitation.

Taken together, our results demonstrate that label-free DDM provides an interpretable, multiscale readout of collective endogenous cytoplasmic fluctuations. Across egg extracts and living cells, the framework resolves how macromolecular composition, energy-driven activity, and cytoskeletal organization shift apparent crowdedness and effective agitation across hierarchical spatiotemporal scales.

## II. RESULTS

### A. Endogenous Cytoplasmic Fluctuations are Subdiffusive and Heterogeneous

As outlined in the introduction, previous studies have identified two striking features of the cytoplasm through the analysis of tracer particle trajectories: viscoelastic subdiffusion and non-Gaussian displacement distributions [39–45]. To determine whether these signatures also manifest in endogenous cytoplasmic fluctuations, we first briefly introduce both concepts within the framework of DDM and then examine label-free bright-field measurements of *Xenopus laevis* egg extract (see *Materials and Methods* and Figure 1A). The theoretical derivations are provided in the *Supplementary Material*.

#### 1. Viscoelastic Subdiffusion

Viscoelastic subdiffusion is commonly described by the fractional Brownian motion (fBm) model and phenomenologically quantified by the anomalous scaling behavior of the (ensemble and/or time-averaged) mean-squared displacement (MSD) as a function of lag time ⟨[Δ*r*(Δ*t*)]⟩^2^ ∝ (Δ*t*)^*α*^, with 0 *< α <* 1 denoting the sub-diffusive regime. Subdiffusive fBm features negatively correlated, stationary increments Δ*r*(Δ*t*) [46, 47], which posits that the ISF is only dependent on the lag time Δ*t*. In the context of DDM, the corresponding ISF is a Gaussian, given by

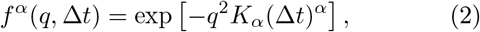

where *K*_*α*_ is the fractional diffusivity in units of m^2^ s^−*α*^. A physical interpretation of the fractional diffusivity is not straightforward because of its dimensionality that depends on the scaling exponent *α*. Therefore, any two fractional diffusivities with different fractional exponents cannot be directly compared; nevertheless, their magnitudes provide important information. To circumvent this, we will consistently report dimensionless diffusivities that are rescaled by their dimensional unit *K*_*α*_*/*µm^2^ s^−*α*^. The fractional exponent *α* may serve as an indicator of ‘molecular crowdedness’ [8, 23, 48, 49] with the lower *α* indicating the more crowded system. Mechanistically, subdiffusive fBm resembles a ‘power-law material’ with viscoelastic properties, captured by the spring-pot model [50] where *α* → 0 indicates a more elastic, ‘spring-like’ response of the system, and *α* → 1 a more viscous, ‘dashpot-like’ behavior. Concordant with Equation (2), for *α* = 1 the system is purely viscous, and the scatterers undergo Brownian motion.

#### 2. Non-Gaussian Displacement Distributions and Superstatistics

The origin of the observed non-Gaussian tails in the van Hove distribution *G*(Δ*r*, Δ*t*) (see e.g., Figure S1) is associated with heterogeneity in the system, specifically a distribution of diffusivities *P* (*K*_*α*_) [44], a concept known as superstatistics [51, 52]. In the context of DDM, this translates to

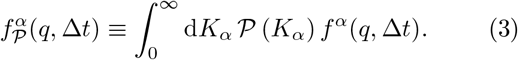

A trivial case occurs when the diffusivity is delta-distributed at a mean value ⟨*K*_*α*_⟩, i.e., *K*_*α*_ ~ *δ*(*K*_*α*_ − ⟨*K*_*α*_⟩)^2^, which yields the ISF

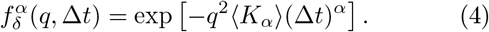

Although this homogeneity assumption fails to account for non-Gaussian tails in the ISF (see Figure S1 and the inset in Figure 1B), it serves as a conservative assumption in the following analysis.

Alternatively, an exponential distribution of diffusivities, *K*_*α*_ ~ ℰ (⟨*K*_*α*_⟩^−1^) (Equation (S12)), as employed in the context of bacterial cytoplasm in the work of [39] and further discussed in [43, 53], leads to

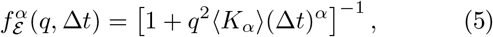

and allows for immobile scatterers (*K*_*α*_ = 0). However, because DDM evaluates changes in intensity over time, very slow or immobilized scatterers contribute only weakly to the signal, which can limit the applicability of Equation (5) in our setting. Additionally, this class of superstatistics might be appropriate for a homogeneous population of tracer particles, but is not necessarily best match for the intrinsically heterogeneous endogenous scatterers in the extract. The notion of a heterogeneous scatterer population in the extract is well motivated by considering the molecular weight distribution of proteins comprising the proteome of *Xenopus laevis* [38]. Besides a strong dependence on the charge, [5] found an inverse relationship between the molecular weight and the diffusivity of various proteins in *Xenopus* egg extract, consistent with the Stokes–Einstein relation [54]. This concept may be further extended to protein complexes and larger aggregates, which all contribute to the measured scattering information.

To describe such a heterogeneous scatterer population in the extract, we propose using an inverse Gaussian (IG) distribution, *K*_*α*_ ~ IG(⟨*K*_*α*_⟩, ⟨*K*_*α*_⟩) (Equation (S15)), which does not include *K*_*α*_ = 0 (immobile scatterers) in its support. This assumption is motivated by a generalized Stokes–Einstein relation hypothesized as *K*_*α*_ ∝ 1*/R*, where *R* is the scatterer radius. The resulting ISF then reads

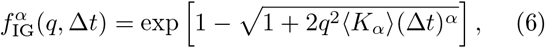

which exhibits non-Gaussian tails, although less pronounced than those in Equation (5) (see *Supplementary Material*; Figure S1). To simplify the model, we assumed in Equation (6) that the mean and standard deviation of the IG distribution are equal (see *Supplementary Material*; Equation (S19)).

It is important to note that various other diffusivity distributions can describe heterogeneous populations. For instance, [55] previously used a gamma-distributed diffusivity (we include the corresponding ISF expression in the *Supplementary Material*; Equation (S22)). While the gamma distribution offers suitable characteristics, its shape and scale parameters are less straightforward to interpret compared to the IG distribution. In addition, one may avoid committing to a specific functional form altogether by using a moment-expansion approximation of Equation (3) [56, 57], which introduces a single ‘relative heterogeneity’ parameter *κ* that quantifies the variance of *P*(*K*_*α*_) relative to its mean (see *Supplementary Material*).

We emphasize that the subsequent procedure is largely unaffected by the choice of either gamma or IG distributions, or moment expansion of Equation (3). However, to maintain consistency with the other model assumptions (specifically having only two open model parameters, ⟨*K*_*α*_⟩ and *α*) we primarily focus on Equation (6) when describing an inherent size distribution of scatterers.

#### 3. Model Fitting and Parameter Extraction

To assess the above theoretical considerations in the context of our model system, we first acquired image sequences of high-speed egg extract (HSS), as described in *Materials and Methods* and schematically illustrated in Figure 1A. The data were analyzed using the DDM algorithm outlined in *Materials and Methods*, yielding ISFs over lag time Δ*t* for each displacement mode *q* (Figure 1B). We fitted the closed-form models Equations (4) to (6) to ℜ [*f* (*q*, Δ*t*)] and extracted ⟨*K*_*α*_⟩ (*q*) and *α*(*q*) (right panel of Figure 1B). Although Equation (6) provided the best fit over Δ*t* based on residual analysis, all three models yielded a strong *q*-dependence of the fitted parameters. Since ⟨*K*_*α*_⟩ and *α* describe the underlying dynamics which should not carry any *q*-dependence, these trends suggest that our diffusivity-based superstatistical approach Equation (3) is further influenced by the optical properties of the heterogeneous scatterer population. To clarify the origin of this, we pursued two complementary strategies. First, we numerically evaluated an effective ISF model that accounts for form-factor weighting in a heterogeneous scatterer population. Second, we performed Monte Carlo simulations as an image-based benchmark of the full acquisition and analysis pipeline.

A minimally extended form of Equation (3), taking the form factor *F* (*q, R*) of each scatterer into account, reads

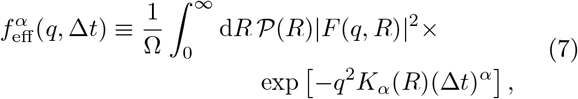

assuming that the complex refractive index and fractional exponent are not correlated with the particle size. In Equation (7), Ω is the normalization constant (see *Supplementary Material*), and *K*_*α*_(*R*) denotes the fractional diffusivity in dependence of the scatterer radius *R*. To link radius and *K*_*α*_(*R*) distributions, we assume that a generalized Stokes–Einstein relation of the form *K*_*α*_ ∝ 1*/R* with a constant ‘fractional’ viscosity *η*_*α*_ in units of Pa s^*α*^ holds. The scatterer radii are then distributed as *R* ~ *k*_B_*T/*6*πη*_*α*_*K*_*α*_ with e.g., *K*_*α*_ ~ IG(⟨*K*_*α*_⟩, ⟨*K*_*α*_⟩), where *T* denotes the temperature. Although these *ad hoc* assumptions are not strictly derived from first principles, they provide a suitable mathematical framework to describe the fBm ISF of a heterogeneous scatterer population in a locally homogeneous fluid. To this end, we evaluated Equation (7), which now contains the effects of the particle form factor, a particle size distribution, and the respective diffusivities numerically. When we fitted this result with our closed-form models Equations (4) to (6), we obtained very similar *q*-dependent trends as in the experimental data (see *Supplementary Material*; Figure S3).

Next, we performed Monte Carlo simulations of particles undergoing fBm and applied the same DDM analysis to the resulting image sequences. In the absence of a particle-size distribution, the *q*-dependence of ⟨*K*_*α*_⟩ and *α* vanished. When we additionally introduced a particle-size distribution (see *Materials and Methods* and *Supplementary Material*; Table S1 and Figures S2A and S3B), the simulations reproduced the qualitative *q*-dependence of *α*(*q*) and ⟨*K*_*α*_⟩ (*q*) observed experimentally. These results show that size polydispersity can generate the fitted *q*-dependent parameter trends even when the underlying dynamics are specified by only the two system parameters *α* and ⟨*K*_*α*_⟩.

We therefore use these *q*-trends resulting from the closed-form fits to define *q*-*independent* best estimates of the system parameters as follows. For the fractional exponent *α*, we use the estimates obtained by fitting the ISF data with Equation (6) for each *q* and define 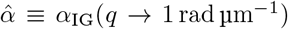 from the lower-limit plateau of a sigmoidal fit over the experimentally accessible *q*-range (see *Supplementary Material*; Equation (S26)).

Similarly, the best estimate of the (dimensionless) fractional diffusivity is obtained by fitting the ISF with Equation (4) and taking the upper-limit plateau of a sigmoidal fit, yielding 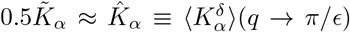, which approximates half the median of the inverse Gaussian distribution 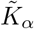 (see *Supplementary Material*). Here, *ϵ* is the empirically determined effective pixel size of the experimental setup (see *Materials and Methods* for further information). The procedure is illustrated in Figure 1B, right (see also the comparison between synthetic simulation inputs and corresponding best estimates presented in the *Supplementary Material*; Figures S2B and S2C).

#### 4. Experimental Results

Applying this framework to a series of eight consecutive measurements (for *N* = 3 samples) depicted in Figure 2 (HSS control), we observe that HSS exhibits stationary behavior, or more accurately, stationarity of the increments, over the observed time window of approximately 130 min, excluding an initial equilibration phase. This initial equilibration phase is presumably due to shear stresses stemming from pipetting and capillary forces employed to float the sample into the microfluidic channel. Excluding the first data point (*t >* 18.5 min), our DDM analysis yields the following long-time-averaged parameter estimates: 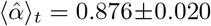, and 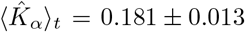, which is in agreement with the notion of cytoplasmic viscoelastic subdiffusion. Additionally, we employed the moment-expanded approximate ISF (Equation (S23)) to obtain an estimate of the relative heterogeneity of the scatterer population *κ* without making explicit assumptions about an underlying diffusivity distribution. We observe that *κ*(*q* ≳ 3.5 rad µm^−1^) ≈ const. (see *Supplementary Material*; Figure S14), and take the median ± 68% confidence interval (CI) over 3.5 rad µm^−1^ ≤ *q* ≤ *π/ϵ* as our best estimate. As shown in Figure 2, 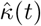 remains approximately constant after the initial equilibration phase of the system. The time-averaged best estimate is given by 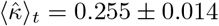.

**FIG. 2:**
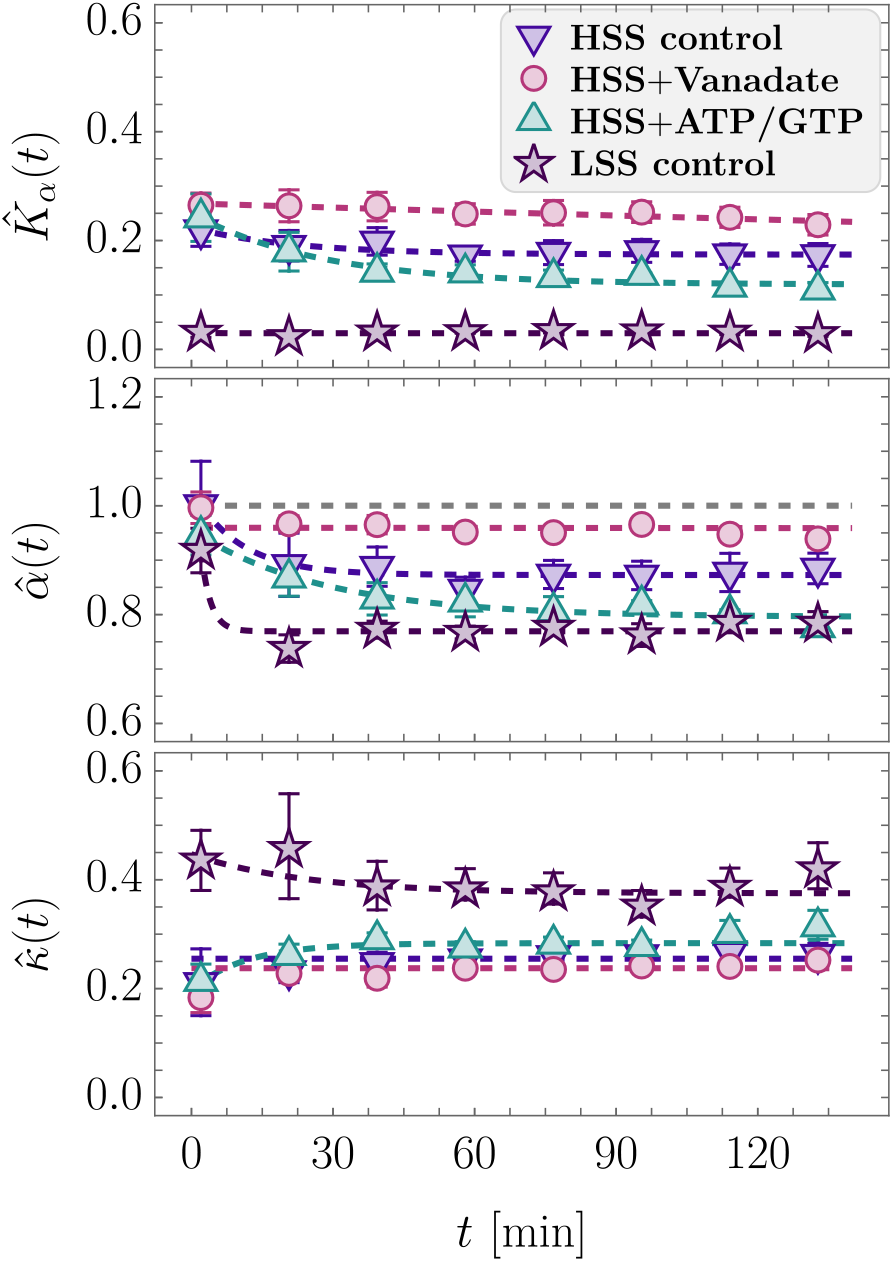
Temporal evolution of the dimensionally normalized fractional diffusivity 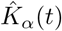 (top), corresponding fractional exponent 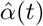 (middle), and relative heterogeneity 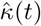 (bottom) for HSS (control; purple down-triangles), HSS + Vanadate (red circles), HSS + 20 × Energy Mix (ATP/GTP; teal up-triangles), and LSS (control; purple stars) as a function of experiment time *t*. Each point displays mean ± pooled SEM from three independent measurements; dashed lines are fits using Equation (8).

To further corroborate the concept of cytoplasmic crowdedness in the context of our model system, we analyzed low-speed *Xenopus laevis* egg extract (LSS) (Figure 1A). Unlike HSS, LSS contains larger macromolecular components, including membranes, ribosomes, and glycogen [58], which are expected to increase the degree of crowdedness. Accordingly, we hypothesized that LSS would exhibit a smaller fractional exponent 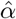 than HSS. Repeating the same measurement protocol as outlined before, we observe that LSS also maintains stationary behavior over the measurement time (Figure 2, LSS control) following an initial equilibration phase. Excluding the first data point, the average parameter best estimates for LSS are 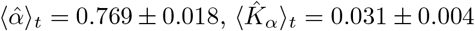, and 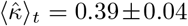. We note that our estimates of the fractional exponents for both HSS and LSS are in good agreement with the values reported in [22], using particle tracking and FCS methods.

These findings indicate a more crowded and heterogeneous environment in LSS compared to HSS, as well as a more elastic-like response, underscoring the influence of macromolecular composition on cytoplasmic dynamics. Thus, the signatures obtained from endogenous fluctuations not only recover the characteristic subdiffusive and heterogeneous behavior of the cytoplasm, but also resolve collective dynamical differences between HSS and LSS in a label-free manner.

### B. Activity-driven Non-stationary Dynamics

Having found HSS and LSS to remain approximately stationary over the measurement period, we next asked whether energy-dependent activity changed this behavior. Stationarity requires that the ISF depends on the lag time Δ*t*, but not on the absolute measurement time *t*; active molecular processes can therefore become apparent through a temporal evolution of the measured dynamics [47, 59, 60]. Non-stationarity may also be associated to aging [61].

To test the effect of energy-dependent activity, we supplemented HSS with ATP and GTP to stimulate processes including polymerization and motor protein activity [2] (see *Materials and Methods*), and referred to this condition as ‘active HSS’. As a complementary control, we inhibited residual ATPase activity by adding Vanadate and referred to this condition as ‘passive HSS’ (see *Materials and Methods*).

Analogously, we performed consecutive DDM measurements over a total duration of approximately 130 min, operating under the assumption that the stationarity-breaking timescale is much larger than a full relaxation of the measured ISF for the accessible displacement modes *q*. The results of these measurements are displayed in Figure 2.

Like the HSS control, the inferred parameters of passive HSS showed little temporal variation over the measurement period. However, passive HSS was consistently less subdiffusive, approaching Brownian-like dynamics, while its relative heterogeneity 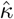 remained approximately constant and slightly below that of the HSS control (see Table I). These differences may partly result from the dilution introduced during Vanadate addition (see *Materials and Methods*), although changes in energy-dependent molecular organization could also contribute. In contrast, active HSS exhibited a progressive decrease in both 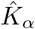 and 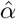, accompanied by a subtle increase in 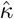, whose final value exceeded those of both passive HSS and the HSS control. Interestingly, 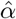 eventually reached values comparable to those of LSS (see Table I).

**TABLE I:** Numerical values from DDM analysis of the cytoplasm: best estimates of the dimensionally normalized fractional diffusivity 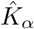, fractional exponent 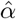, and relative heterogeneity 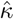 under indicated conditions, averaged across three independent repetitions. For HSS+ATP/GTP, the final time series value is reported (see Figure 2); for other cases, long-time averages (excluding the first data point) and uncertainty estimates are provided.

| Condition | $\hat{K}_\alpha$ | $\hat{\alpha}$ | $\hat{\kappa}$ |
| --- | --- | --- | --- |
| **HSS Control** | $0.181 \pm 0.013$ | $0.876 \pm 0.020$ | $0.255 \pm 0.014$ |
| **HSS+Vanadate** | $0.250 \pm 0.015$ | $0.955 \pm 0.012$ | $0.236 \pm 0.012$ |
| **HSS+ATP/GTP** | $0.110 \pm 0.013$ | $0.776 \pm 0.013$ | $0.314^{+0.030}_{-0.023}$ |
| **LSS Control** | $0.031 \pm 0.004$ | $0.769 \pm 0.018$ | $0.39 \pm 0.04$ |

To characterize this temporal evolution, we assumed that active HSS transitions from an initial state 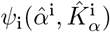 to a final state 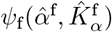 with characteristic timescale *τ*_*ψ*_ and fitted the time series with a single-exponential relaxation

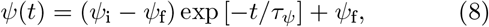

to determine the characteristic timescales 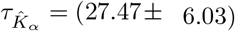 min and 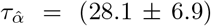 min, which coincide well within the margins of uncertainty. The similarity of these timescales suggests that the changes in 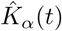 and 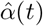 may arise from a common underlying non-equilibrium remodeling process. This activity-driven evolution shifts the system toward slower, more subdiffusive, and more heterogeneous dynamics, consistent with activity-induced heterogeneity and a more elastic-like response.

### C. Microtubule Networks Mediate Dynamical Hierarchies

Having examined changes in cytoplasmic dynamics driven by composition and activity, we next asked how a cytoskeletal network alters these dynamics. We stabilized microtubules (MTs) by adding Taxol to HSS at a concentration of *c*_Tax_ = 0.25 µm, optimized to minimize global contractions [62, 63] while preserving MT network formation (see *Materials and Methods* and *Supplementary Material*). Strikingly, the resulting ISF exhibited a second characteristic timescale (Figure 3), qualitatively distinct from the parameter shifts observed for composition and activity.

**FIG. 3:**
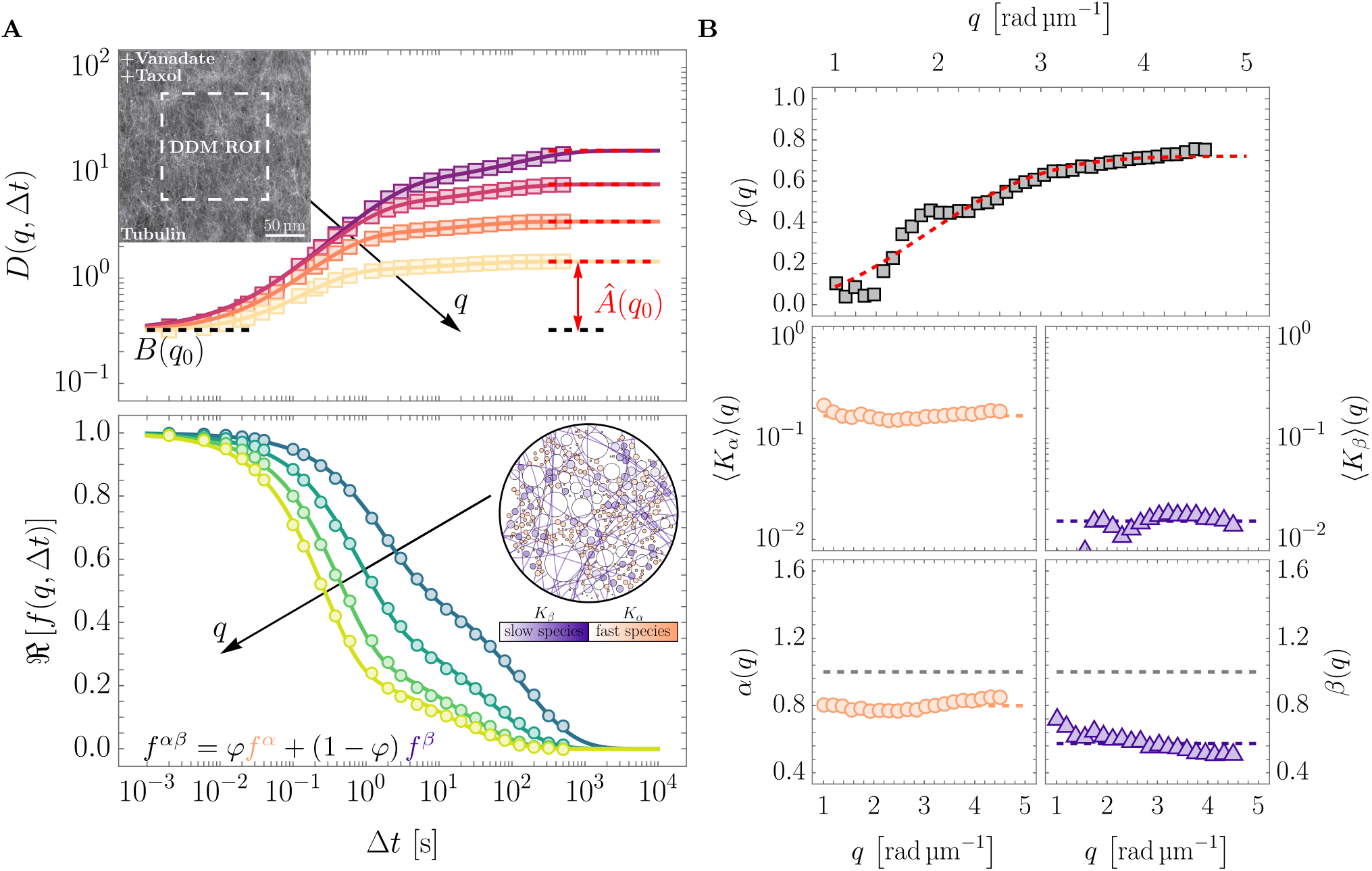
Stabilized MT networks induce hierarchical heterogeneities in the passive HSS. **A** Top: Azimuthally averaged DDM map *D*(*q*, Δ*t*) with black arrow indicating dispersion over *q*, black dashed line showing camera noise *B*(*q*_0_), and red double arrow indicating amplitude *Â*(*q*_0_) for *q*_0_ ≈ 4.13 rad µm^−1^; inset: fluorescence image of HyLyte 488-labeled MT network (ROI marked). **A** Bottom: Experimental ISF ℜ [*f* (*q*, Δ*t*)] (circles) and fits with Equation (9) (solid lines). Inset: Schematic of the freeze-in scenario via the MT network. **B**: *q*-dependence of fitted parameters; top: *φ*(*q*) with stretched sigmoidal fit Equation (10) (red dashed); middle: fractional mean diffusivities ⟨*K*_*α*_⟩ (*q*) and ⟨*K*_*β*_⟩ (*q*) (dashed: medians); bottom: exponents *α*(*q*) and *β*(*q*), with medians (colored dashed lines), and *α* = *β* = 1 for reference (gray dashed lines).

To understand this behavior phenomenologically, we described the data using an effective two-state fBm model

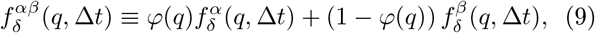

where *φ*(*q*) represents the displacement-mode-dependent relative contribution of the two states (see Figure 3A inset). We interpret the two contributions as a slower *β*-state associated with the MT network and a faster *α*-state associated with cytosolic dynamics. Similar two-state descriptions have been employed for heterogeneous dynamics in two-dimensional glassy colloids [64] and bidisperse colloidal suspensions [65], while a general theoretical treatment of Markovian switching between two states was presented in [66]. In our context, Equation (9) can be interpreted as a ‘frozen mixture’, in which the contributing entities do not switch between states. We hypothesize that the *β*-state comprises scattering contributions from the MT network together with sufficiently large particles coupled to or captured by the network, whereas the *α*-state predominantly reflects smaller particles that remain comparatively unaffected by the mesh [67].

Fitting Equation (9) to the experimental ISF, as shown in Figure 3A, and applying suitable constraints, we obtained five fit parameters as functions of the displacement mode *q*. We find that the mean fractional diffusivities ⟨*K*_*α*_⟩ and ⟨*K*_*β*_⟩ and the corresponding fractional exponents *α* and *β* remain approximately constant across the available *q* range. Consistent with the previous analysis, we therefore take half of the median ± 68% CI for the fractional diffusivities, and the median ± 68% CI for the fractional exponents over *q* as best estimates. Conversely, *φ*(*q*) displays a dependence on *q* (see Figure 3B).

Building on [64], where a similar ISF model was employed, we modeled *φ*(*q*) using a stretched exponential function

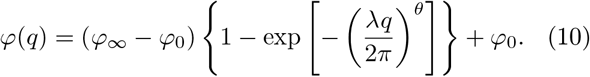

Note that we utilize a stretched exponential with variable exponent *θ*, whereas [64] used a fixed value *θ* = 2. Fitting Equation (10) yielded the parameter best estimates 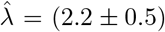 µm and 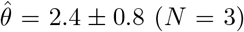. The inferred values of *φ*_0_ and *φ*_∞_ showed significant variability between experiments. We interpret *λ* as a characteristic displacement length scale marking the crossover between the two dynamical contributions: above this scale, the measured ISF is increasingly dominated by the slower *β*-state, whereas below it the faster cytosolic *α*-state contributes more strongly.

The two states exhibit clearly distinct dynamical parameters. The slow *β*-state is characterized by 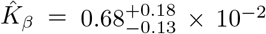 and 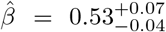, compared with 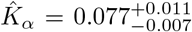 and 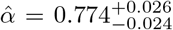 for the faster *α*-state. Interestingly, the exponent of the *β*-state is consistent with the *β* = 1*/*2 scaling characteristic of Rouse-like relaxation in polymer chains [68]. Its strongly subdiffusive character is also consistent with fractional exponents previously reported for tracer dynamics in reconstituted actin–MT networks and for MT bending fluctuations [69, 70]. The *α*-state itself is also slower and more subdiffusive than the corresponding passive HSS control 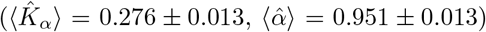, indicating that MT assembly not only introduces a distinct slow dynamical contribution but also alters the background cytosolic dynamics.

To test whether a size-selective interaction with the MT network can account for the observed two-timescale ISF, we conducted Monte Carlo simulations of a freeze-in model, as elaborated in the *Supplementary Material*. In brief, starting from the inverse Gaussian distribution of diffusivities introduced above, we hypothesized that the MT network captures larger scatterers and imposes new dynamical properties on them, while smaller scatterers remain unaffected. We implemented this by introducing a cutoff probability *p*, which splits the scatterer distribution into two subpopulations: those with *K* ≥ *K*^∗^ assigned to the *α*-state and those with *K < K*^∗^ assigned to the *β*-state. Under the assumed generalized Stokes– Einstein relation between diffusivity and scatterer size, this effectively corresponds to a separation into two size groups. The results of these simulations are shown in the *Supplementary Material*; Figures S4 and S5. They demonstrate that our DDM analysis recovers the input parameters within uncertainty over a range of *p* values, while also reproducing the experimentally observed dependence of *φ*(*q*).

Interestingly, we can choose a value 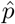 for which the simulated crossover length 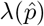 matches the experimentally measured 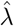 (see Figure S5). This fixes the corresponding cutoff value *K*^∗^ and, through the generalized Stokes– Einstein relation, yields a model-predicted characteristic scatterer radius at the crossover between the *α*-and *β*-states, 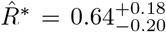 µm. We then wondered whether this theoretical prediction corresponds to an independently measurable structural length scale of the MT network. To this end, we segmented fluorescence images of the MT network and obtained a pore-radius distribution with mode 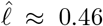 µm (see *Supplementary Material*; Figure S9), in remarkable agreement with the predicted crossover radius 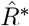. The combined dynamical, simulation, and structural evidence supports the interpretation that the secondary timescale in the measured ISF arises from MT-mediated modification of the dynamic scattering contributions of larger cytoplasmic scatterers.

### D. Microtubules Agitate and Crowd the Cytoplasm of Living Cells

Having established a label-free readout of cytoplasmic dynamics in egg extract, we next asked whether the same approach could be extended to living cells, where composition, activity, and cytoskeletal organization coexist within a confined cellular environment. We therefore applied DDM to phase-contrast recordings of individual HeLa cells. To obtain sufficient endogenous contrast at the single-cell level, we used phase-contrast microscopy with halogen illumination, which also provided improved optical sectioning for cell-scale measurements. We focused on adherent HeLa cells, which remained sufficiently stationary over the duration of a measurement (~ 20 min; see *Supplementary Material*; Figure S11). We acquired large fields of view (≈207.6 µm × 207.6 µm) and segmented individual cells using Cellpose [71]. A representative field of view and an example single-cell region of interest are shown in Figures 4A and 4B.

**FIG. 4:**
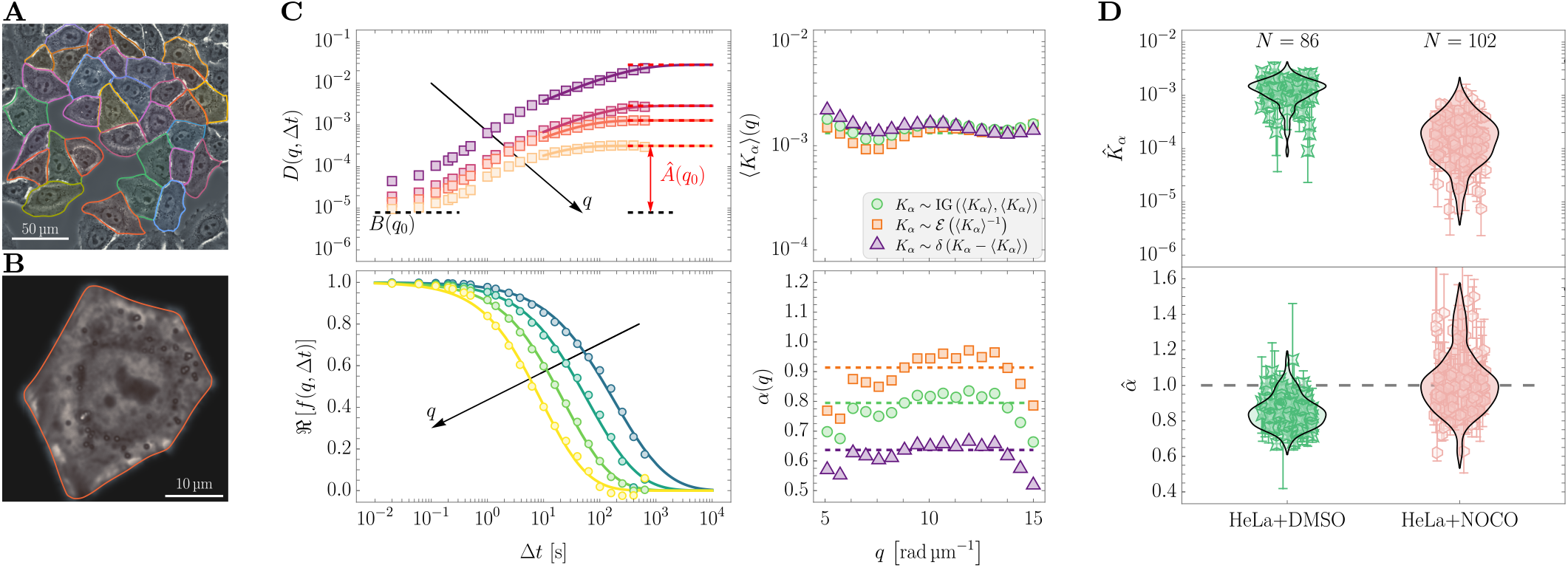
DDM analysis of HeLa cells reveals subdiffusive intracellular dynamics depend on the microtubule network. **A**: Representative phase-contrast field of view with segmented cells outlined. **B**: Example single-cell region of interest used for DDM analysis. **C**: DDM analysis for the cell depicted in **B** showing the structure function *D*(*q*, Δ*t*) with fitted amplitude *Â*(*q*) and background *B*(*q*), the corresponding ℜ [*f* (*q*, Δ*t*)], and the fitted parameter trends *K*_*α*_(*q*) and *α*(*q*) over the analysis range in analogy to Figure 1B. **D**: Summary of best-estimate parameters 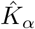 and 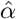 across cells for control (HeLa+DMSO) and Nocodazole-treated (HeLa+NOCO) conditions. Points show individual cells (median ± 68% CI) and the violin outlines indicate the distributions.

Applying the DDM pipeline to each segmented cell yielded single-cell ISFs, as shown in Figure 4C. For the initial cell-level analysis, we considered the whole cell, including both cytoplasm and nucleus. In contrast to the passivated and MT-stabilized egg extract, where a second decay was visible within the observation window, the single-cell HeLa ISFs showed a single dominant decay over the accessible (*q*, Δ*t*) range. We fitted the three closed-form models (Equations (4) to (6)) to the single-cell ISFs. Across models, the extracted ⟨*K*_*α*_⟩ (*q*) trends were comparable, while *α*(*q*) showed model-dependent dispersion, as depicted in Figure 4C. For the cell-level estimates reported here, we selected the inverse Gaussian model based on the fitting residuals and defined, for each cell, the median ± 68% CI of the fitted parameter values over the analysis range as best estimates. For control cells (HeLa+DMSO, *N* = 86), we obtained 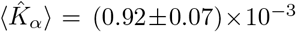 and 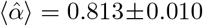 (weighted averages), indicating subdiffusive dynamics that were substantially slower than in egg extract. Compartment-resolved analysis of control cells yielded comparable fractional exponents across whole-cell, nuclear, and cytoplasmic regions, while diffusivities were lower in the nucleus and higher in the cytoplasm relative to the whole-cell average (see *Supplementary Material*; Figure S10).

To probe how MT perturbations affect these dynamics, we treated cells with Nocodazole at a concentration of 1 µm, which was insufficient to fully depolymerize the network (see *Materials and Methods* and Figure S12). Repeating the same acquisition, segmentation, and fitting procedure, we observed a pronounced reduction in the fractional diffusivity, while the fractional exponent shifted toward more Brownian-like, and in some cases even superdiffusive (*α >* 1), behavior. For Nocodazole-treated cells (HeLa+NOCO, *N* = 102), we obtained 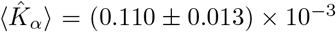 and 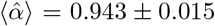 (weighted averages).

This observation suggests that the dynamic MTs in living cells do not only increase the effective crowdedness (lower fractional exponent), in line with our extract results, but also agitate the apparent dynamics (higher diffusivities).

## III. DISCUSSION AND OUTLOOK

Our results show that endogenous cytoplasmic fluctuations provide quantitative signatures of molecular crowdedness, heterogeneity, activity, and cytoskeletal organization. Across egg extract and living cells, these signatures reveal how distinct physical perturbations reshape the spatiotemporal dynamics of the cytoplasm, giving rise to the notion of hierarchical heterogeneities.

We found that the DDM signal in egg extract is consistent with a heterogeneous population of endogenous scatterers. Within a phenomenological description, closed-form subdiffusive models capture the lag-time dependence well, but exhibit systematic *q*-dependent effective parameters. Both the heterogeneous, form-factor-weighted ISF model and the image-based Monte Carlo simulations reproduce these trends when analyzed with the same closed-form models, supporting that the observed *q*-dependence can arise from heterogeneity of the scattering population. The extracted best estimates should therefore be interpreted as effective parameters of the dominant endogenous fluctuations contributing to the optical signal. A more complete microscopic description would require explicit knowledge of the optical contrast and physicochemical properties of the contributing macromolecular species, which remains difficult for a system as complex as cytoplasm. The present framework instead provides a consistent label-free basis for comparing dynamical states across conditions.

Time-resolved measurements further separated passive stationarity from activity-driven evolution of the system. HSS and LSS controls, as well as passivated HSS, remained approximately stationary over the time of measurment, whereas ATP/GTP addition drove a progressive decrease in both 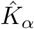 and 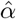, together with a subtle increase in 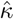. In contrast, Vanadate-treated HSS exhibited almost Brownian-like dynamics, although both perturbations involved some dilution of the extract. The close agreement between the characteristic timescales of 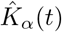 and 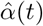, both around 30 min, suggests that these changes are linked to a common underlying non-equilibrium remodeling process. Interestingly, this observation is consistent with the findings of [22], who reported activity-dependent slowing down of particles larger than approximately 100 nm in *Xenopus* egg extract. We therefore view the ATP/GTP condition as an active state that drives the extract toward slower, more subdiffusive, and more heterogeneous dynamics, consistent with increased effective crowdedness and altered macromolecular organization [12, 22]. Such changes in crowdedness may in turn influence downstream biochemical processes [4] and could be related to activity-driven self-assembly and the formation of mesoscale structures akin to protocells previously observed in *Xenopus* egg extract [6, 10, 22].

Stabilizing MTs with Taxol in the passivated extract introduced an additional hierarchy in the ISF. In this condition, we observed a second decay and described the data with a two-state fBm mixture model. To our knowledge, a second characteristic subdiffusive timescale mediated by Taxol-stabilized MT networks in *Xenopus* egg extract has not been reported previously. The analysis separates a faster cytosolic *α*-state from a slower, more sub-diffusive *β*-state, with the latter exhibiting a fractional exponent close to the *β* = 1*/*2 scaling characteristic of Rouse-like dynamics in polymer chains [68]. Its strongly subdiffusive character is also consistent with fractional exponents previously reported for tracer dynamics in reconstituted actin–MT networks and for MT bending fluctuations [69, 70]. We further identified a characteristic crossover length scale 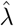, beyond which the measured ISF is increasingly dominated by the slow component. In our simplest interpretation, the *β*-state reflects fluctuations associated with the MT network together with larger scatterers coupled to or trapped by it, while the *α*-state predominantly reflects smaller-scale cytosolic fluctuations. Consistent with this picture, the freeze-in simulations reproduced the observed *q*-dependent mixing, and the inferred crossover radius 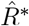 showed remarkable agreement with the independently measured MT pore radius.

The experiments further indicated that the cytosolic *α*-state itself was slower and more subdiffusive relative to the control, suggesting that MT assembly also alters the effective cytosolic environment. One possible mechanism is that the MT meshwork, together with larger scatterers coupled to or trapped by it, forms an effective mesh that also constrains the faster cytosolic component.

Measurements in HeLa cells showed that a similar phenomenology can be extracted in living matter. The single-cell ISFs showed one dominant decay within the accessible window and did not resolve a clear second relaxation process as observed in the MT-stabilized extract. A possible explanation is that the MT-associated contribution to the optical signal is considerably weaker in living cells than in the dense Taxol-stabilized meshwork of the extract, such that a corresponding relaxation does not emerge as a distinct second decay. Control cells (HeLa+DMSO) exhibited subdiffusive dynamics, whereas Nocodazole treatment reduced 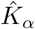, in line with the findings of [72], and shifted 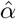 toward more Brownian-like values. One interpretation consistent with our extract results is that MTs contribute two distinct dynamical effects in cells. A polymerized MT network can increase effective crowdedness and mechanical constraints, promoting subdiffusion, while dynamic MTs and associated motor activity can introduce agitation and re-arrangements that increase the effective diffusivity. Partial MT perturbation with Nocodazole can therefore reduce this agitation component, yielding slower yet more Brownian-like effective dynamics detected with DDM.

We summarized the parameter space explored in this study in Figure 5. Interestingly, compositional variation between HSS and LSS, energy addition to HSS, and MT assembly in passivated HSS yield similar fractional exponents around 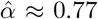 despite distinct fractional diffusivities. These exponents are furthermore concordant with the fractional scaling observed in control HeLa cells, although the corresponding fractional diffusivity in cells is substantially lower. This recurrence raises the possibility that cytoplasmic systems tend toward a characteristic range of effective crowdedness while retaining substantial variation in the speed of endogenous fluctuations. Whether this reflects a preferred dynamical regime or distinct mechanisms producing similar scaling remains to be established.

**FIG. 5:**
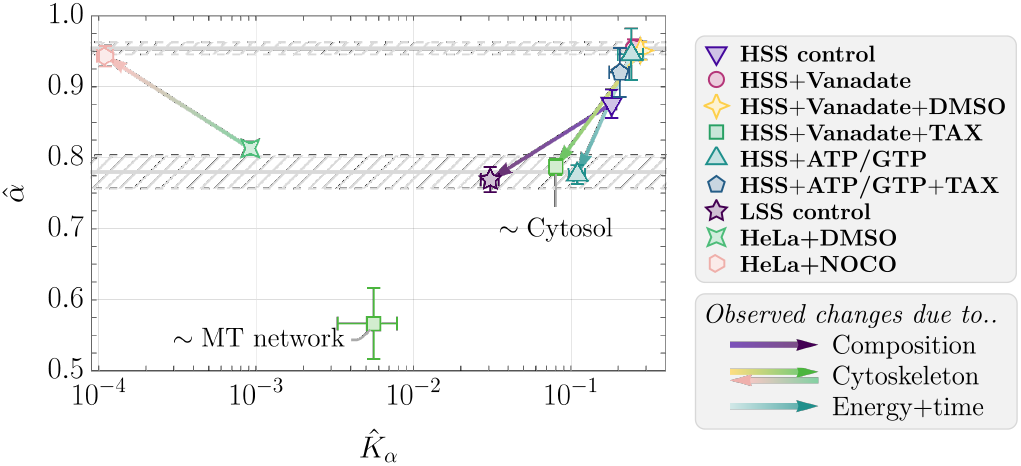
Overview of the parameter space (best estimates of the dimensionally normalized fractional diffusivity 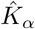 and fractional exponent 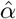) of *Xenopus laevis* egg extract and HeLa cells explored in this study. All experiments except HSS+ATP/GTP represent long-time averages with uncertainty estimates, excluding the first data point (see *Materials and Methods* and main text for details). For HSS+ATP/GTP, initial and final values from time-series measurements are shown (symbols), with the arrow indicating the temporal progression. Gray hatched areas and lines represent constant fit estimates of 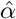 for the cases HSS+ATP/GTP, HSS+Vanadate, HSS+Vanadate+DMSO (top), and HSS+ATP/GTP, LSS control, HSS+Vanadate+Taxol (bottom), respectively, with 68% CIs. For HSS+ATP/GTP+Taxol, data were acquired outside the contracting MT band (see *Supplementary Material*; Figure S8). HeLa data (HeLa+DMSO and HeLa+NOCO) are population estimates from single-cell DDM (see Figure 4).

The comparison to previous measurements is consistent with this picture. Particle-tracking and FCS studies in *Xenopus* egg extract have reported subdiffusive dynamics of tracer particles and proteins [22, 67, 73]. Interestingly, the Gaussian displacement distributions reported for a comparatively homogeneous probe population in [73] are consistent with our interpretation that the non-Gaussianity observed here arises primarily from the heterogeneity of the endogenous scattering population rather than from the dynamics of any single species.

Looking ahead, our DDM-based framework could be used to examine how endogenous cytoplasmic fluctuation signatures change under additional perturbations of living cells, for example during osmotic shocks [74], senescence-associated remodeling [75], or disease-relevant perturbations [13]. Beyond stationarity, out-of-equilibrium concepts such as mean-back relaxation [76, 77] may provide complementary probe-free measures of active processes directly from intensity fluctuations.

On the modeling side, beyond the superstatistical arguments invoked here, our results could be further interpreted in the context of more fundamental established theories, such as fBm with random scaling exponents [78, 79], multifractional Brownian motion [80], diffusing diffusivities [44, 81, 82], and subordination processes [53], but may also give rise to new theoretical connections. These approaches hold great potential to better understand the physical mechanisms underlying the highly complex nature of cytoplasmic diffusion. Improving forward simulations to include more realistic scatterer ensembles and optical contrast mechanisms, including aggregation, interaction-driven signatures, and refractive-index variability, would further help connect the fitted effective parameters to microscopic constituents and facilitate their biological interpretation in the complex cytoplasm.

## IV. MATERIALS AND METHODS

### A. High-Speed *Xenopus laevis* Egg Extract Production

Low-speed (LSS) and high-speed (HSS) extracts were prepared as described previously [83, 84]. Briefly, freshly laid *Xenopus laevis* eggs were collected, washed in physiological buffers, and subjected to a crushing spin at 19,000 × *g* for 20 min. The crude cytoplasmic layer (LSS) was collected, driven into interphase with CaCl_2_ (final concentration 0.6 mm), and supplemented with cycloheximide (final concentration 0.1 mg mL^−1^) and cytochalasin D (final concentration 0.01 mg mL^−1^). To generate HSS, the crude LSS was centrifuged at 260,000 × *g* for 90 min at 4°C to remove dense components (membranes, glycogen, ribosomes, mitochondria). The cytosolic layer was collected and spun again at 260,000 × *g* for 30 min at 4°C. Final HSS aliquots were snap frozen in liquid nitrogen and stored at −70°C.

### B. CSF-XB Buffer

CSF-XB buffer contained 10 mm HEPES, 50 mm sucrose, 100 mm KCl, 1 mm MgCl_2_, 0.1 mm CaCl_2_ (from XB-salt solution), and 5 mm EGTA. The pH was adjusted to 7.7 with KOH. The buffer was filtered (200 nm), aliquoted, and stored at 4°C.

### C. 20*×* XB-Salt Solution

A 20× XB-salt solution contained 2 mol L^−1^ KCl, 20 mm MgCl_2_, and 2 mm CaCl_2_. The solution was filtered (200 nm), aliquoted, and stored at 4°C.

### D. 20*×* Energy Mix

The 20× Energy Mix contained 100 mm creatine phosphate, 0.5 mg L^−1^ creatine kinase, 5 mm ATP, and 5 mm GTP. Specifically, 131.6 µg ATP and 100 µg GTP were dissolved in 1 mL double-distilled water and combined with 571.4 µg creatine phosphate dissolved in 1 mL double-distilled water. Creatine kinase was prepared by diluting 5 µL of a 0.5 mg L^−1^ stock with 95 µL double-distilled water and added to the mixture. Finally, 3.1 µL of 1 mol L^−1^ MgCl_2_ stock and 2 µL of 1.5 mol L^−1^ DTT were added. Aliquots were stored at −70°C.

### E. Cell culture

HeLa Kyoto cells were maintained in high glucose + GlutaMAX DMEM (Thermo Fisher Scientific 61965026) supplemented with 10% heat-inactivated fetal bovine serum (Sigma Aldrich F7524-500ML) and 1% penicillin-streptomycin (Thermo Fisher Scientific 15070063). Cells were grown in 25 cm^2^ flasks (Greiner Bio-One 690175) at 37°C and 5% CO_2_ and passaged every 2 days at 80% confluency (1:10). Detachment was performed with 0.25% Trypsin-EDTA (Thermo Fisher Scientific 25200072). Cell counts and viability were assessed using a LUNA II™ automated cell counter (BioCat L40001-LG) with Erythrosin B (BioCat L13002-LG).

### F. Sample Preparation and Mounting

*Xenopus* extracts (HSS or LSS; 30 µL) were thawed, briefly centrifuged, and kept on ice to suppress microtubule (MT) formation. Drug stocks (in CSF-XB or DMSO) were kept on ice and added immediately before loading. For experiments without MT stabilization, 3 µL of 5.5 mm Vanadate (Sigma S6508) in CSF-XB or 3 µL of 20× Energy Mix were added. For MT stabilization, 1.5 µL of 11 mm Vanadate in CSF-XB was added first, followed by 1.5 µL of 5.5 µm Taxol (Sigma 33069-62-4) in DMSO (Sigma 5.89569). MT labeling was performed by adding HyLyte 488-labeled tubulin (Cytoskeleton, Inc. TL670M-A) to the respective Vanadate or Energy Mix condition as described in the datasheet. Samples were gently mixed, loaded into Ibidi µ-Slide VI 0.4 channels, sealed with lukewarm VALAP (1:1:1 vaseline:lanolin:paraffin), transferred to the microscope, and measured within 2 min. For HeLa experiments, cells (1.7 × 10^6^ mL^−1^) were seeded into Ibidi µ-Slide I 0.4 channels and incubated at 37°C for at least 24 h. Channels were washed four times with medium containing Nocodazole in DMSO (Sigma 31430-18-9) at 1 µm f.c. or with DMSO at matched dilution and incubated for 1 h before imaging.

### G. Experimental Setup and Measurement Scheme

*Xenopus* egg extract measurements were performed on an inverted bright-field microscope (Zeiss Axio Observer 3) with a quasi-monochromatic LED (Luminus CBT-120, (623 ± 19) nm), a 20× /0.45 objective (Zeiss N-Achroplan), and a CMOS camera (Mikrotron EoSens). 8-bit images (200 px × 200 px) were acquired at 2 ms exposure, with focus near the lower channel surface to reduce axial diffusion contributions. HeLa measurements were performed on an Olympus IX72 stand with a 40× /0.65 PH2 objective (Olympus PLN) and internal 1.6× magnification, using a halogen lamp (Osram) and a Hamamatsu ORCA FUSION camera. 16-bit images (2048 px × 2048 px) were acquired at 2 ms exposure. Camera control and acquisition were implemented in `Python` using `pylablib` [85]. Image acquisition was performed in three phases: 5000 frames at 500 Hz (*Xenopus* only), then 5000 frames at 50 Hz, and 5000 frames at 5 Hz. Data and metadata were stored in `hdf5` format. For HeLa analysis, cells were segmented with Cellpose [71], cropped to 400 px × 400 px around the centroid, and multiplied by Gaussian-smoothed masks to reduce contributions from neighboring cells. Effective pixel sizes were determined with a USAF test target (Thorlabs) as *ϵ*_BF_ ≈ 0.68 µm px^−1^ and *ϵ*_PC_ ≈ 0.10 µm px^−1^ for bright-field and phase-contrast modes, respectively. Displacement modes were computed as *q*_BF_ ≈ {0.0459, …, 4.5945} rad µm^−1^ and *q*_PC_ ≈ {0.155, …, 30.92} rad µm^−1^, where *q* = 2*π/*(*hϵ*), …, *π/ϵ* and *h* is the image size. Modes *q*_BF_ *<* 1 rad µm^−1^ and *q*_PC_ *<* 5 rad µm^−1^ as well as *q*_PC_ *>* 15 rad µm^−1^ were excluded due to parasitic contributions, lower statistics, incomplete relaxation, or insufficient signal-to-noise ratio. Fluorescence data were acquired on a Zeiss LSM 980 with AiryScan 2. *Xenopus* samples were measured at *T* = (21±1)°C and HeLa samples at *T* ≈ 37°C using a heated stage insert (PeCon).

### H. Simulation Scheme

Simulations were performed using custom `Mathematica` [86] scripts. For each scenario, trajectories of *N* = 2 × 10^3^ particles ***s***_*j*_(*t*) in 2D with periodic boundary conditions were generated by `RandomFunction` for the respective stochastic process. Intensities were rendered on a 200 px × 200 px grid as *I*_obj_(***x***, *t*) = ⟨*I*_0_⟩ *T* (***x***, *t*), where ⟨*I*_0_⟩ is the background intensity and *T* (***x***, *t*) is the sample transmittance. For simplicity, scatterers were modeled as overlapping opaque disks with

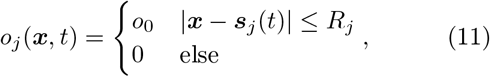

where 0 ≤ *o*_0_ ≤ 1 and ***s***_*j*_(*t*) denotes the particle center. Radii were assigned via 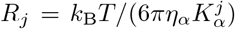 µm, with 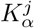 sampled from *P*(*K*_*α*_) (see *Supplementary Material*), setting *η*_*α*_ = 1 mPa s^*α*^. Transmittance was constructed using the `Opacity` compositing rule in `Mathematica` [86]. For three overlapping particles

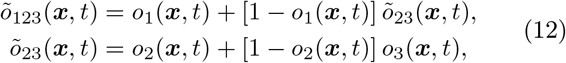

with *õ*_23_(***x***, *t*) = *õ*_32_(***x***, *t*). For *N* particles, *õ*_1…*N*_ (***x***, *t*) = *o*_1_(***x***, *t*) + [1 − *o*_1_(***x***, *t*)] *õ*_2…*N*_ (***x***, *t*), and *T* (***x***, *t*) = 1 − *õ*_1…*N*_ (***x***, *t*).

Images were generated as

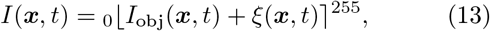

emulating 8-bit rounding and clipping. The additive noise term *ξ*(***x***, *t*) was sampled independently for each pixel and frame from a Poisson distribution with mean *µ*_0_. Thus, ⟨*ξ*⟩ = Var(*ξ*) = *µ*_0_, with *µ*_0_ controlling the noise strength. For each simulation, sequences of 1000 images at different frame rates were generated using *o*_0_ = 0.5.

### I. Differential Dynamic Microscopy

DDM analysis was performed using custom `Mathematica` [86] scripts following [29] and the main text. Images were multiplied by a Blackman-Harris window [87], differences were computed over lag times, and the corresponding Fourier transforms were azimuthally averaged to obtain *D*(*q*, Δ*t*) via Equation (1). Data from different frame-rate blocks were stitched by element-wise averaging over a minimal lag-time overlap. For experiments without MT stabilization, only 500 Hz and 50 Hz blocks were used. For MT-stabilized extracts, the 5 Hz block was included. For HeLa, 50 Hz and 5 Hz blocks were used.

For *Xenopus* measurements, *B*(*q*) was determined from no-sample background measurements by constructing and interpolating a background map *D*_0_(⟨*I*_0_⟩, *q*, Δ*t*) over mean intensity and taking *B*(*q*) = *D*_0_(⟨*I*⟩, *q*, Δ*t*)_Δ*t*_ at the measurement intensity ⟨*I*⟩. We adjusted all measurements to ⟨*I*⟩ ≈ 139 to operate in a regime where the noise level is largely independent of intensity fluctuations (see *Supplementary Material*; Figure S13). For HeLa, *B*(*q*) was estimated from *D*(*q*, Δ*t* = 0.02 s). For all experiments, the amplitude *Â*(*q*) was estimated by fitting a (double) sigmoidal amplitude model *g*(*q*, Δ*t*) to *D*(*q*, Δ*t*) and taking *Â*(*q*) = *g*(*q*, Δ*t* → ∞) − *B*(*q*) (see Figure 1B and Equation (S26)), with initial values *Â*_0_(*q*) = 2⟨|*I*(*q, t*)|^2^⟩_*t*_ − *B*(*q*). For MT-stabilized experiments, we imposed *A*(*q*) ≤ 2 |⟨*I*(*q, t*)|^2^⟩_*t*_ − *B*(*q*). The ISF was then obtained by numerically solving Equation (1) and fitted with the theoretical model of choice. Goodness of fit was assessed from residuals averaged across lag time for each displacement mode.

### J. Statistical Analysis

All statistical analyses were performed using custom `Mathematica` [86] scripts. Data fitting and extraction of the best fit estimates of a parameter *p* along with the corresponding confidence interval (CI) estimates were achieved using the `NonlinearModelFit` function. The pooled SEM, associated with the (weighted) average over *N* experimental repetitions 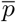, was computed as

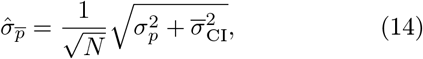

where *σ*_*p*_ is the standard deviation of *p*, and 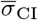 indicates the average of the estimated CIs. Weighted averages and corresponding standard deviations were computed using the `NonlinearModelFit` function, specifying the individual weights as 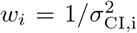. The uncertainty estimates for time averages were calculated as

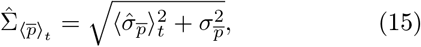

with the time-averaged pooled SEM 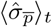 and the temporal standard deviation of 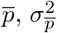, over *N*_*t*_ time points.

## Author Contributions

C.M., V.Z. and J.G. conceptualized the project. C.M. developed the methodology, and conducted the experiments. All authors interpreted the experimental results. C.M., and V.Z. conducted the theoretical considerations. All authors interpreted the theoretical results. C.M. wrote the original draft of this manuscript. All authors edited and improved the manuscript. A.B., and S.R. provided the *Xenopus laevis* egg extract. C.S. performed cell culture. J.G., and V.Z. supervised the project.

## Acknowledgments

The authors thank Manuela Hauke for the preparation of buffers, the TDSU4 Lab-on-a-chip systems for usage of the bright-field microscope, and Olga Lyraki, Jana Bachir-Salvador, Tomohisa Toda, Kyoohyun Kim, Shada Abuhattum, Timon Beck, and Sara Kaliman for discussions.

## Conflicts of Interest

The authors declare no competing interests.

## Data Availability Statement

The data and code that support the findings of this study can be accessed under the following link https://doi.org/10.17617/3.FT6V60.

## Footnotes

1 The 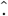 notation indicates best estimates throughout this study.

2 The ~ notation reads “distributed as”.

## Notes

### Competing Interest Statement

The authors have declared no competing interest.

### Summary of Updates

Added living cell data in a new Figure 4 and new section, improved theoretical model and overall clarity of the manuscript.

