## Supplementary Materials for "Hierarchical Heterogeneities in Spatiotemporal Dynamics of the Cytoplasm"


### Derivations for DDM

In the following, we recapitulate how the DDM observable $D(q, \Delta t)$ is connected to the normalized intermediate scattering function (ISF).

#### The Intermediate Scattering Function

Following the derivations in [1, 2], here we consider the dynamical properties in space and time of a given sample via the number density $\rho(\boldsymbol{x}, t)$, where $\boldsymbol{x} = (x, y)^{\mathrm{T}}$ denotes the 2D Cartesian coordinates in the lateral plane. Assuming that the sample consists of $N$ discrete scatterers, the number, or one-point density and associated density modes are given by

$$\begin{aligned} \rho(\boldsymbol{x}, t) &\equiv \sum_{j=1}^{N} \langle \delta\left[\boldsymbol{x} - \boldsymbol{s}_j(t)\right]\rangle, \text{ and} \\ \rho(\boldsymbol{k}, t) &= \sum_{j=1}^{N} \langle \exp\left[-i\boldsymbol{k} \cdot \boldsymbol{s}_j(t)\right]\rangle, \end{aligned} \tag{S1}$$

respectively, with $\boldsymbol{s}_j(t)$ being the position of particle $j$ at time $t$, and $\langle \dots \rangle$ denoting the ensemble average. Further assuming that the $N$ scatterers undergo processes with stationary increments, we introduce the time-averaged, lag-time–dependent van Hove correlation function, or normalized two-point density, as

$$\begin{aligned} G(\Delta\boldsymbol{r}, \Delta t) &\equiv N^{-1} \sum_{i=1}^{N}\sum_{j=1}^{N} \langle \delta\left[\Delta\boldsymbol{r} - \boldsymbol{s}_i(t+\Delta t) + \boldsymbol{s}_j(t)\right]\rangle_t \\ &\equiv G_{\mathrm{s}}(\Delta\boldsymbol{r}, \Delta t) + G_{\mathrm{d}}(\Delta\boldsymbol{r}, \Delta t), \end{aligned} \tag{S2}$$

where $\langle \dots \rangle_t$ denotes time and ensemble averaging in this instance. $G_{\mathrm{s}}(\Delta\boldsymbol{r}, \Delta t)$ is the so-called self-part, the probability density function of the displacements of particle $i$ at time $t + \Delta t$ with respect to itself at time $t$, and $G_{\mathrm{d}}(\Delta\boldsymbol{r}, \Delta t)$ is the distinct part, the probability density function of displacements of particle $j \neq i$ at time $t + \Delta t$ with respect to particle $i$ at time $t$. The spatial Fourier transform of the van Hove correlation function is called the intermediate scattering function (ISF), and can be expressed as

$$F(\boldsymbol{q}, \Delta t) \equiv N^{-1} \langle \rho(-\boldsymbol{q}, t)\rho(\boldsymbol{q}, t + \Delta t)\rangle_t. \tag{S3}$$

Note that in Equation (S3)$\langle \dots \rangle_t$ indicates the time average. We may further define the time-averaged static structure factor $S(\boldsymbol{q}) \equiv N^{-1}\langle \rho(-\boldsymbol{q}, t)\rho(\boldsymbol{q}, t)\rangle_t$, and it is customary to define the normalized time-averaged, lag-time–dependent ISF as

$$\begin{aligned} f(\boldsymbol{q}, \Delta t) &\equiv \langle F(\boldsymbol{q}, \Delta t)\rangle_t / S(\boldsymbol{q}) \\ &= f_{\mathrm{s}}(\boldsymbol{q}, \Delta t) + f_{\mathrm{d}}(\boldsymbol{q}, \Delta t), \end{aligned} \tag{S4}$$

which is referred to as the ISF throughout this study.

#### The DDM Observable

As motivated earlier, instead of considering number densities $\rho(\boldsymbol{x}, t)$, (bright-field) DDM employs intensity maps of spatially extended objects. In the focal plane, the recorded image modes may be written as

$$I(\boldsymbol{x}, t) = \mathcal{T}(\boldsymbol{x}) \circledast I_{\mathrm{obj}}(\boldsymbol{x}, t) + \xi(\boldsymbol{x}, t), \tag{S5}$$

where $\mathcal{T}(\boldsymbol{x})$ is the (effective) point-spread function in the focal plane, $\circledast$ denotes spatial convolution, and $\xi(\boldsymbol{x}, t)$ denotes camera noise. In our experiments, digital images are acquired by a CMOS sensor under quasi-monochromatic Köhler illumination. In this study, the relevant contrast arises from endogenous, weakly scattering objects embedded in a predominantly transmitting medium. We therefore adopt a linear, space-invariant image formation description for the in-plane modes, based on the van Cittert–Zernike theorem (see e.g., [3]). For an optically homogeneous population of scatterers with in-plane positions $\boldsymbol{x}_j(t)$ and characteristic size $R_j$, the Fourier modes of the image intensity can be expressed as

$$I(\boldsymbol{q}, t) \propto \mathcal{T}(\boldsymbol{q}) \sum_j F(\boldsymbol{q}, R_j) \exp\left[i\boldsymbol{q} \cdot \boldsymbol{x}_j(t)\right] + \xi(\boldsymbol{q}, t), \tag{S6}$$

where $\mathcal{T}(\boldsymbol{q})$ is the effective optical transfer function, including finite axial sectioning, and $F(\boldsymbol{q}, R)$ is the corresponding effective particle form factor [4]. In the present description, we introduce $F(\boldsymbol{q}, R)$ as a section-averaged quantity that results from weighting the phase-delay and absorption contrast along the optical axis $z$ by the microscope-specific optical sectioning function. It therefore describes how an object of radius $R$ contributes, on average, to the image mode $\boldsymbol{q}$ in the focal plane. (S6) is the starting point for the DDM observable in the presence of polydispersity. The experimentally relevant consequence is that the measured dynamics are optically weighted by $|F(\boldsymbol{q}, R)|^2$. Following the standard DDM construction [5, 6, 7], we consider image differences at lag time $\Delta t$ and define

$$\begin{aligned} D(\boldsymbol{q}, \Delta t) &\equiv \left\langle |I(\boldsymbol{q}, t+\Delta t) - I(\boldsymbol{q}, t)|^2 \right\rangle_t \\ &\equiv \hat{A}(\boldsymbol{q}) \left\{1 - \Re\left[f(\boldsymbol{q}, \Delta t)\right]\right\} + B(\boldsymbol{q}), \end{aligned} \tag{S7}$$

where $B(\boldsymbol{q})$ is the noise floor and $\Re[f(\boldsymbol{q}, \Delta t)]$ is the real part of the normalized intermediate scattering function (ISF) associated with the contrast mechanism of the images. For delta-correlated camera noise, $\langle \xi(\boldsymbol{x}, t)\xi(\boldsymbol{x}, t+\Delta t)\rangle = \hat{\sigma}_0^2(\boldsymbol{x})\delta(\Delta t)$, one has $B(\boldsymbol{q}) = 2\sigma_0^2(\boldsymbol{q})$. The amplitude term $\hat{A}(\boldsymbol{q}) \equiv 2\langle |I(\boldsymbol{q}, t)|^2\rangle_t - B(\boldsymbol{q})$ follows from the optical transfer and the static structure factor of the image contrast. In the present context, (S6) yields the usual factorization

$$\hat{A}(\boldsymbol{q}) \propto |\mathcal{T}(\boldsymbol{q})|^2 S(\boldsymbol{q}), \tag{S8}$$

with $S(\boldsymbol{q})$ the (contrast-weighted) static structure factor. For isotropic dynamics, we will use the azimuthal average $q = |\boldsymbol{q}|$. We now make explicit how polydispersity enters the dynamic term. Consider a population with size distribution $\mathcal{P}(R)$ and single-particle dynamics that, conditional on $R$, exhibit a fBm-like ISF

$$f(q, \Delta t; R) = \exp\left[-q^2 K_\alpha(R)(\Delta t)^\alpha\right]. \tag{S9}$$

Here, $K_\alpha(R)$ is the generalized transport coefficient or diffusivity. For the special case of Brownian dynamics ($\alpha = 1$), $K_1(R) = D(R)$ and, in the Stokes–Einstein limit

$$K_\alpha(R) = \frac{k_B T}{6\pi\eta_\alpha R}, \tag{S10}$$

with $\eta_\alpha$ the generalized viscosity parameter. Inserting (S6) into the DDM correlation construction (S7), and averaging over a polydisperse population, the measured ISF becomes a form-factor-weighted mixture

$$f_{\mathrm{eff}}^{\alpha}(q, \Delta t) = \frac{\int_0^\infty \mathrm{d}R\, \mathcal{P}(R)\, |F(q, R)|^2 \exp\left[-q^2 K_\alpha(R)(\Delta t)^\alpha\right]}{\int_0^\infty \mathrm{d}R\, \mathcal{P}(R)\, |F(q, R)|^2}. \tag{S11}$$

(S11) shows that systematic $q$-dependence of parameters obtained by fitting closed-form ISFs that average only over $K_\alpha$ can arise even when the underlying dynamics are consistent across $q$. With a change in $q$ the effective weighting over $R$ through $|F(q, R)|^2$ is changed, and therefore the apparent ISF.

Additional contrast heterogeneities can be incorporated analogously. For example, if each object carries an additional multiplicative contrast factor (e.g. refractive-index contrast) that is narrowly distributed and uncorrelated with $R$, it rescales the amplitude $\hat{A}(q)$ but leaves $f_{\mathrm{eff}}^{\alpha}(q, \Delta t)$ unchanged. Correlations between contrast and size would introduce an additional weighting factor in (S11). In the main text and in the analysis of samples without microtubule stabilization, we treat these effects as subleading and retain the effective description used in Eq. (3) of the main text.

### Super-statistical Models

In this appendix, we introduce theoretical ISF models based on super-statistical arguments, following the ideas and results of [8, 9, 10]. We focus on distributions of (fractional) diffusivities $\mathcal{P}(K_\alpha)$ and their consequences for the van Hove function and the ISF. The main conclusions are summarized in Figure S1. In addition, we use these results to motivate and validate the parameter estimation scheme used in the main text, and to compare analytical fits to numerical evaluation of an effective ISF and to Monte Carlo simulations (see Figure S3 and Figure S2).

#### Fractional Brownian Motion with a Distribution of Diffusivities

##### Exponential Distribution of Diffusivities

Following [8], in complex biological matter one may assume that the diffusivity of similarly sized scatterers depends on position due to local changes in crowdedness. In this scenario, some tracers could be effectively immobile, others might be slowed down, while another fraction moves freely. A continuous distribution that captures this notion is the exponential distribution with PDF

$$\mathcal{E}(x \geq 0; \lambda > 0) \equiv \lambda \exp\left(-\lambda x\right), \tag{S12}$$

where $x$ is the ordinate and $\lambda$ is the characteristic parameter. Using Eq. (3) in the main text with Equation (S12), one obtains a Laplace distribution of displacements

$$G_{\mathcal{E}}^{\alpha}(\Delta r, \Delta t) = \sqrt{\frac{\lambda}{4\left(\Delta t\right)^\alpha}} \exp\left[-\Delta r \sqrt{\frac{\lambda}{\left(\Delta t\right)^\alpha}}\right], \tag{S13}$$

which is the PDF of $\mathcal{L}\left(\Delta r; \mu = 0, \sigma = \sqrt{\lambda^{-1}\left(\Delta t\right)^\alpha}\right)$. The corresponding ISF is

$$f_{\mathcal{E}}^{\alpha}(q, \Delta t) = \frac{1}{1 + q^2 \lambda^{-1} \left(\Delta t\right)^\alpha}. \tag{S14}$$

For $\lambda = \langle K_\alpha \rangle^{-1}$ we recover Eq. (5) of the main text.

#### Inverse Gaussian Distribution of Diffusivities

In the context of DDM, inhomogeneities can also arise from a heterogeneous scatterer population, for instance via a distribution of scatterer sizes. For the Brownian case ($\alpha = 1$), assuming a constant viscosity and the Stokes–Einstein relation, an exponential distribution of diffusivities is not physical in the limit $K \to 0 \Leftrightarrow R \to \infty$, since $\mathcal{E}(K = 0; \lambda = 1/\langle K \rangle) = 1/\langle K \rangle$ remains finite. A distribution that suppresses weight at $K \to 0$ is the inverse Gaussian distribution with PDF

$$\mathrm{IG}(x > 0; \mu, \sigma) \equiv \sqrt{\frac{\sigma}{2\pi x^3}} \exp\left[-\frac{\sigma (x-\mu)^2}{2x\mu^2}\right], \tag{S15}$$

and mode

$$\hat{\mu} = -\frac{3\mu^2}{2\sigma} + \frac{\sqrt{9\mu^4 + 4\mu^2\sigma^2}}{2\sigma}. \tag{S16}$$

The inverse Gaussian distribution arises naturally from a first passage time problem of a Brownian motion with drift. Following the approach above, the displacement distribution reads

$$G_{\mathrm{IG}}(\Delta r, \Delta t) = \frac{\sigma \exp(\sigma/\mu)}{\pi\mu\sqrt{\sigma\Delta t + (\Delta r)^2}} \mathcal{K}_1\left[\frac{1}{\mu\Delta t}\sqrt{\sigma\Delta t + (\Delta r)^2}\right], \tag{S17}$$

where $\mathcal{K}_n(z)$ is the modified Bessel function of the second kind, satisfying $z^2 y'' + z y' - (z^2 + n^2) y = 0$ [11]. Fourier transformation of Equation (S17) yields

$$f_{\mathrm{IG}}(q, \Delta t) = \exp\left[\sigma/\mu - \sqrt{\sigma^2/\mu^2 + 2q^2\sigma\Delta t}\right]. \tag{S18}$$

For $\mu = \sigma$, Equation (S18) reduces to

$$f_{\mathrm{IG}}(q, \Delta t) = \exp\left[1 - \sqrt{1 + 2q^2\mu\Delta t}\right]. \tag{S19}$$

At this point, the argument about a size distribution holds only for a purely viscous and sufficiently dilute system. Accounting for viscoelastic effects in terms of fBm is not straightforward, since there is no general relationship between the friction term $\gamma_\alpha$ and the particle radius $R$ for the generalized Stokes–Einstein relation $K_\alpha = k_{\mathrm{B}}T/\gamma_\alpha m$. The relationship between $K_\alpha$ and $R$ depends on system properties. In a minimal attempt to connect a size distribution to viscoelastic subdiffusion, we employ Equation (S10). This yields

$$f^{\alpha}_{\mathrm{IG}}(q, \Delta t) = \exp\left[1 - \sqrt{1 + 2q^2\langle K_\alpha\rangle (\Delta t)^\alpha}\right], \tag{S20}$$

which we use in the main text to connect non-Gaussianity to a size distribution.

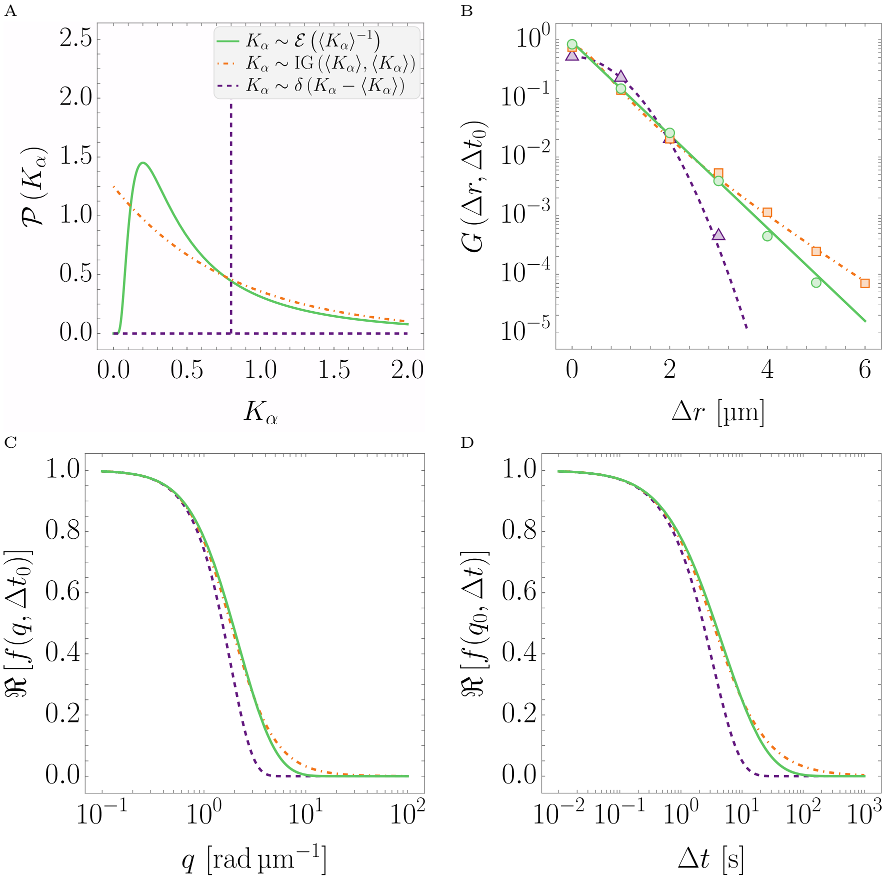


Figure S1: Results of superstatistical analysis of fBm for different diffusivity distributions with mean $\langle K_\alpha\rangle$ and fractional exponent $\alpha$. A PDFs of diffusivity distributions $\mathcal{P}(K_\alpha)$: exponential distribution Equation (S12) (orange, dot-dashed), inverse Gaussian distribution Equation (S15) (green, solid), and delta distribution (purple, dashed). B Displacement distributions at fixed lag time for the cases shown in A. Markers indicate MC simulation results and lines are analytical results. C and D Real part of the ISF for the cases shown in A as a function of lag time $\Delta t$ and displacement mode $q$, respectively.

#### Gamma Distribution of Diffusivities

Analogously, the superstatistical ISF for a gamma distribution reads as follows. The PDF is

$$\gamma(x > 0; \mu, \nu) = \frac{\nu^{-\mu}}{\Gamma(\mu)} \exp(-x/\nu)\, x^{-1+\mu}, \tag{S21}$$

where $\mu$ and $\nu$ are the shape and scale parameters, and $\Gamma(\dots)$ is the gamma function. The corresponding ISF is

$$f_\gamma^\alpha(q, \Delta t) = \left[1 + q^2 \nu\, (\Delta t)^\alpha\right]^{-\mu}. \tag{S22}$$

For $\mu = 1$ and $\nu = \lambda^{-1}$ we recover the Laplacian case Equation (S14).

#### Moment Expansion of the superstatistical ISF

Adding to this discussion, the dynamics of polydisperse protein-rich clusters in the context of DDM have been studied in [12]. There, the authors expanded the ISF in Eq. (3) in the main text (for $\alpha = 1$) around the mean diffusivity $K_\alpha - \langle K_\alpha \rangle$ up to second order. Assuming that the first and second moments of the diffusivity distribution exist, this yields the approximate non-Gaussian ISF [13]

$$f_2^\alpha(q, \Delta t) = f_\delta^\alpha(q, \Delta t) \left[1 + \frac{\mu_2}{2} q^4 (\Delta t)^{2\alpha}\right], \tag{S23}$$

where the 'relative heterogeneity' can be expressed as $\kappa \equiv \mu_2 / \langle K_\alpha \rangle^2$, with

$$\mu_2 \equiv \int_0^\infty \mathrm{d}K_\alpha\, \mathcal{P}(K_\alpha) \left(\langle K_\alpha \rangle - K_\alpha\right)^2. \tag{S24}$$

In practice, $\mu_2$ (or equivalently $\kappa$) may be treated as an open parameter that captures the second moment of $\mathcal{P}(K_\alpha)$ without committing to a specific functional form. An exemplary result for $\kappa$ in dependence of $q$ is shown in Figure S14 for HSS.

### Monte Carlo Simulations of Superstatistical Scenarios

The results of the DDM analysis of MC simulations of the superstatistical diffusion scenarios described above are depicted in Figure S2 and Table S1. The simulations generate trajectories of a heterogeneous particle population with assigned diffusivities (and, where applicable, sizes), render image sequences, and apply the same DDM analysis as used for experimental data. A comparison with the fitting results $\langle K_\alpha \rangle(q)$, $\alpha(q)$, and fit residuals of the experiments, and with numerical calculations using Equation (S11), is shown in Figure S3.

### Numerical Evaluation of $f_{\mathrm{eff}}^\alpha(q, \Delta t)$

For numerical evaluation of Equation (S11), we use the same $(q, \Delta t)$ grid as in the experiments and the MC simulations and choose a phenomenological form factor of the form

$$F(q, R) \propto \frac{R^2}{1 + (qR)^2/2}, \tag{S25}$$

which captures suppression at large $qR$ and emphasizes larger scatterers at fixed $q$. The numerical results and the comparison to experimental trends are shown in Figure S3 for the case of $R \sim k_\mathrm{B}T/6\pi\eta_\alpha K_\alpha$, with $K_\alpha \sim \mathrm{IG}(K_\alpha; \langle K_\alpha \rangle, \langle K_\alpha \rangle)$, and $\eta_\alpha = 1\,\mathrm{mPa\,s}^\alpha$.

### Parameter Estimation

The numerical evaluation of Equation (S11) and the image-based MC simulations both show that fitting the closed-form expressions Eqs. (4) to (6) of the main text to the resulting ISFs induces a characteristic $q$-dependence of fitted parameters. This motivates the estimation scheme used in the main text. In brief, we define $\hat{\alpha}$ from the low-$q$ limit of $\alpha_{\mathrm{IG}}(q)$ obtained from a sigmoidal fit of the type

$$g(q) = (g_\infty - g_0)[1 - f(q)] + g_0 \tag{S26}$$

as $\hat{\alpha} \equiv \alpha_{\mathrm{IG}}(q \to 1\,\mathrm{rad\,\mu m}^{-1})$, where $g_\infty \equiv \lim_{q\to\infty} g(q)$, $g_0 \equiv \lim_{q\to 0^+} g(q)$, and $f(q) = \exp(-aq)$, where $a > 0$. We define $\hat{K}_\alpha$ from the high-$q$ limit of $\langle K_\alpha \rangle_\delta(q)$ obtained from a sigmoidal fit, $\hat{K}_\alpha \equiv \langle K_\alpha^\delta \rangle(q \to \pi/\epsilon)$. The systematic comparison between simulation inputs and these best estimates for the parameter sets used in this work is provided in Figure S3 and Table S1.

### Two-State Processes

Following the derivations of [14, 15], we may express a two-state process that consists of two fBm processes with average diffusivities $\langle K_\alpha \rangle, \langle K_\beta \rangle$ and fractional exponents $\alpha, \beta$ as

$$\begin{aligned} f_\delta^{\alpha\beta}(q, \Delta t) =& \varphi(q) \exp\left[-q^2 \langle K_\alpha \rangle (\Delta t)^\alpha\right] + \\ & (1 - \varphi(q)) \exp\left[-q^2 \langle K_\beta \rangle (\Delta t)^\beta\right], \end{aligned} \tag{S27}$$

where $\varphi(q)$ is the occupation or volume fraction of particles that follow the $\alpha$ process, and, consistently, $1 - \varphi(q)$ is the occupation or volume fraction of particles that follow the $\beta$ process.

Table S1: Overview of the different numerical simulation schemes performed in this study (see *Materials and Methods* and the main text for further information). In cases **a)**–**c)**, the scatterers follow a fractional diffusivity distribution $\mathcal{P}(K_\alpha)$ while their radii $R$ are the same. For cases **d)**–**i)**, the scatterers follow a fractional diffusivity distribution $\mathcal{P}(K_\alpha)$ and their radii are calculated using a generalized Stokes–Einstein relation Equation (S10), with $T = 23^\circ$C and a constant 'fractional' viscosity $\eta_\alpha = 1\,\mathrm{mPa\,s}^\alpha$. The corresponding results of the DDM analysis are displayed in Figure S2.

| | $\alpha$ | $\mathcal{P}(K_\alpha)$ | $\langle K_\alpha \rangle$ | $R$ [µm] |
|---|---|---|---|---|
| **a)** | 0.8 | $\delta(K_\alpha - \langle K_\alpha\rangle)$ | 0.8 | 0.44 |
| **b)** | 0.8 | $\mathcal{E}\left(\langle K_\alpha\rangle^{-1}\right)$ | 0.8 | 0.44 |
| **c)** | 0.8 | $\mathrm{IG}(\langle K_\alpha\rangle, \langle K_\alpha\rangle)$ | 0.8 | 0.44 |
| **d)** | 0.8 | $\mathrm{IG}(\langle K_\alpha\rangle, \langle K_\alpha\rangle)$ | 0.8 | $\sim 1/K_\alpha$ |
| **e)** | 0.6 | $\mathrm{IG}(\langle K_\alpha\rangle, \langle K_\alpha\rangle)$ | 0.8 | $\sim 1/K_\alpha$ |
| **f)** | 0.9 | $\mathrm{IG}(\langle K_\alpha\rangle, \langle K_\alpha\rangle)$ | 0.8 | $\sim 1/K_\alpha$ |
| **g)** | 0.95 | $\mathrm{IG}(\langle K_\alpha\rangle, \langle K_\alpha\rangle)$ | 0.5 | $\sim 1/K_\alpha$ |
| **h)** | 0.95 | $\mathrm{IG}(\langle K_\alpha\rangle, \langle K_\alpha\rangle)$ | 0.3 | $\sim 1/K_\alpha$ |
| **i)** | 0.95 | $\mathrm{IG}(\langle K_\alpha\rangle, \langle K_\alpha\rangle)$ | 1.0 | $\sim 1/K_\alpha$ |

To examine the applicability of Equation (S27), we consider the following freeze-in scenario. In this scenario, we employ an IG distribution of diffusivities $K \sim \mathrm{IG}(\langle K\rangle, \langle K\rangle)$ with a corresponding size of scatterers, as described above, and define a cut-off $K^*$, such that

$$\int_0^{K^*} \mathrm{d}K\, \mathrm{IG}(K; \langle K\rangle, \langle K\rangle) = p. \tag{S28}$$

This cut-off defines two diffusivity populations: $K \to K_\alpha$ for $K \geq K^*$, i.e., smaller scatterers, and $K \to K_\beta$ for $K < K^*$, i.e., larger scatterers. The freeze-in mechanism is then captured by slowing the diffusivity of the $\beta$-process down by tenfold, $K_\beta \to K_\beta/10$, and assigning $\beta \to 0.6$, whereas for the $\alpha$-process, no changes occur. In this model, $p$ effectively tunes the length scale of the freeze-in mechanism.

The results of the DDM analysis of the freeze-in scenario MC simulations described above are depicted in Figure S4 and Figure S5.

#### Effect of Axial Decorrelation

Under partially coherent illumination, the detected image contrast is weighted by an optical sectioning function around the focal plane rather than collected uniformly over the full sample depth. Axial diffusion therefore contributes an additional decorrelation term when the characteristic axial displacement becomes comparable to the sectioning width [7]. We describe this contribution by

$$f_z(\Delta t) = \left[1 + \frac{K_{\alpha,z}(\Delta t)^\alpha}{w_z^2}\right]^{-1/2}, \tag{S29}$$

where $K_{\alpha,z}$ is the generalized transport coefficient in the axial direction and $w_z$ is the effective optical sectioning width. The observed ISF is then approximated by

$$f_{\mathrm{obs}}(q, \Delta t) = f_{\parallel}(q, \Delta t) f_z(\Delta t), \tag{S30}$$

where $f_{\parallel}(q, \Delta t)$ denotes the in-plane ISF. Axial decorrelation becomes appreciable only when $K_{\alpha,z}(\Delta t)^\alpha / w_z^2$ approaches unity and is expected to be most visible at low $q$, where the in-plane relaxation is slow.

For the quasi-monochromatic LED illumination used in the extract experiments, we estimate $w_z \simeq \lambda^2/\Delta\lambda \approx 20\,\mu\mathrm{m}$. To obtain a conservative upper estimate of the axial contribution, we set $K_{\alpha,z} = K_\alpha$. As shown in Figure S6, for the displacement modes considered in the manuscript ($q \geq 1\,\mathrm{rad}\,\mu\mathrm{m}^{-1}$), the in-plane ISF is already largely decayed before axial decorrelation becomes substantial. Including Equation (S29) in the numerical evaluation produces only minor changes in the fitted parameters and residuals. Moreover, axial motion accelerates the observed decorrelation and would therefore bias the analysis toward faster apparent dynamics, tending to reduce rather than generate the slow relaxation and enhanced heterogeneity reported here.

### Microtubule stabilization in *Xenopus laevis* egg extract

To probe how a cytoskeletal component reshapes the DDM-detectable fluctuation spectrum, we stabilized MTs in high-speed *Xenopus laevis* egg extract (HSS) by adding Taxol. This induces the formation of a network within the microfluidic channel that undergoes bulk contraction [16, 17] (see Figure S8). [16, 17] demonstrated that this contraction is correlated with dynein activity and Taxol concentration. While these contractions are noteworthy, they typically occur on much larger length scales ($\propto 1\,\mathrm{mm}$) than our experimental field of view ($\approx 136\,\mu\mathrm{m} \times 136\,\mu\mathrm{m}$), and the direction and magnitude of MT motion depend on the position within the channel. This spatial variability complicates extracting meaningful information using our DDM configuration. To mitigate contractile motion, we attempted to passivate HSS by adding Vanadate. However, Vanadate did not fully inhibit contraction, consistent with observations in [16] involving additional inhibition of dynein. We found that contractions were minimized at low Taxol concentrations. Therefore, to reduce contractions within the observation time while maintaining an MT network, we selected $c_{\mathrm{Tax}} = 0.25\,\mu\mathrm{M}$ as optimal for investigating network-associated fluctuations. We note that this concentration is lower than in [16, 17], which cover $1\,\mu\mathrm{M} \leq c_{\mathrm{Tax}} \leq 25\,\mu\mathrm{M}$.

Figure S7 summarizes time-series DDM measurements for the MT-stabilized condition (HSS+Vanadate+Taxol) and the corresponding control (HSS+Vanadate+DMSO) using the two-state model. While most inferred parameters remain approximately constant over the measurement duration, we observe clear aging in the slow-state transport coefficient $\hat{K}_\beta(t)$,

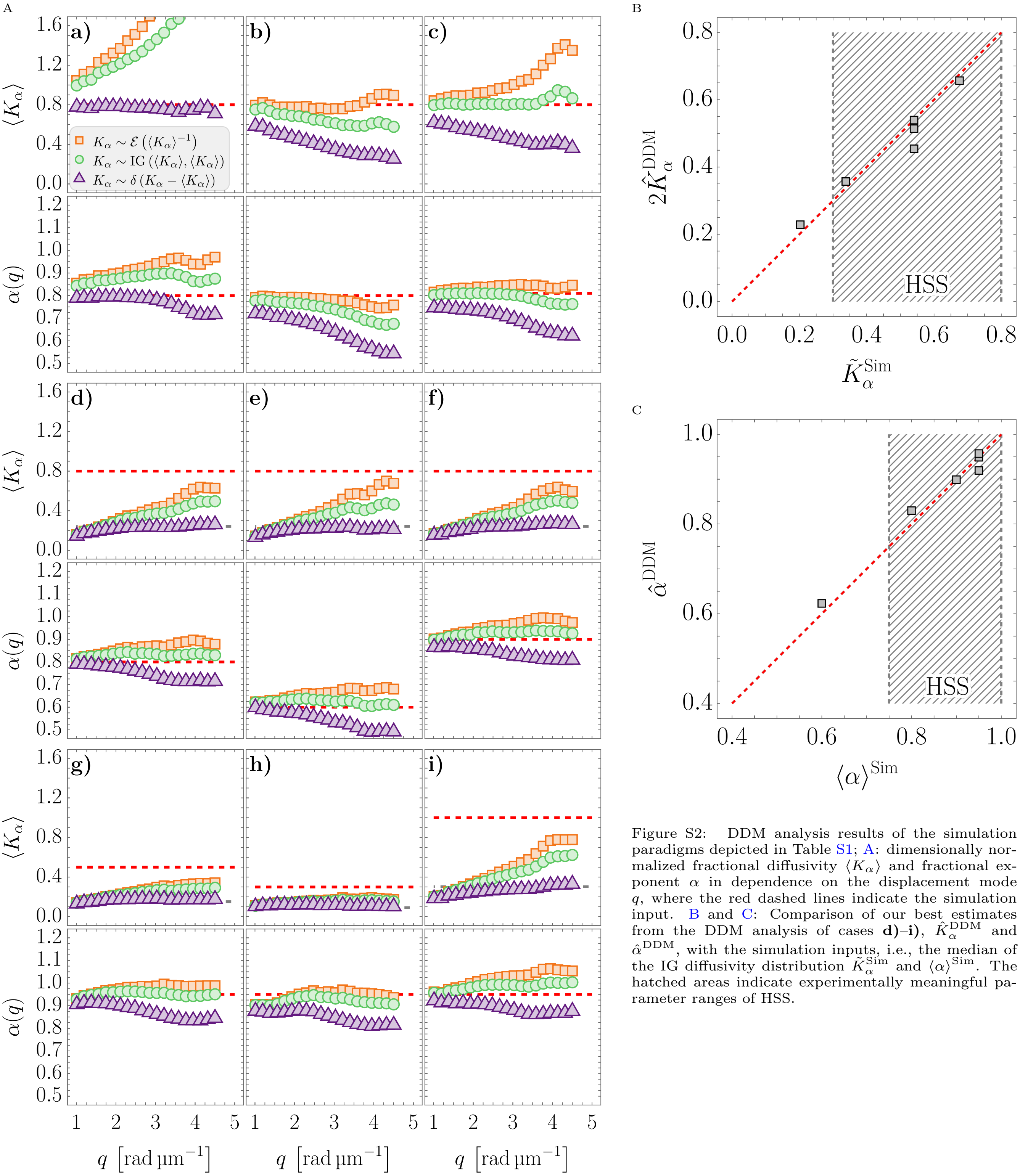


Figure S2: DDM analysis results of the simulation paradigms depicted in Table S1; A: dimensionally normalized fractional diffusivity $\langle K_\alpha \rangle$ and fractional exponent $\alpha$ in dependence on the displacement mode $q$, where the red dashed lines indicate the simulation input. B and C: Comparison of our best estimates from the DDM analysis of cases **d)**–**i)**, $\hat{K}_\alpha^{\mathrm{DDM}}$ and $\hat{\alpha}^{\mathrm{DDM}}$, with the simulation inputs, i.e., the median of the IG diffusivity distribution $\tilde{K}_\alpha^{\mathrm{Sim}}$ and $\langle \alpha \rangle^{\mathrm{Sim}}$. The hatched areas indicate experimentally meaningful parameter ranges of HSS.

and a weaker aging trend in the slow-state exponent $\hat{\beta}(t)$. The origin of this aging effect remains speculative but might be linked to the formation of longer and more stable MT assemblies. By contrast, $\hat{\alpha}(t)$ and $\hat{K}_\alpha(t)$ show no pronounced drift within the uncertainty, and the mixing function parameters remain stable. The characteristic length scale $\hat{\lambda}(t)$ and the stretching exponent $\hat{\theta}(t)$ are consistent with constant values over time.

Figure S8 shows an exemplary picture of a typical microfluidic channel employed for the DDM measurements. Here, MTs were stabilized using Taxol and ATP/GTP was added to allow motor-driven contraction of the MT network. A contraction band is clearly visible at the channel scale, and the same effect can be observed under both fluorescence and bright-field imaging. Importantly, DDM analysis performed outside the contraction band yields an ISF with a single dominant relaxation process, as shown in Figure S8B. In addition, when we allowed the MT network to contract by adding ATP and GTP to the system and chose a higher Taxol concentration ($c_{\mathrm{Tax}} = 5\,\mu\mathrm{M}$) to amplify the effect, measurements outside the MT contraction band did not exhibit a double decay in the ISF (see Fig. S8). This confirms that the secondary decay quantified above is attributable to the presence of the MT network and reinforces its role in mediating dynamical hierarchies within the cytoplasm.

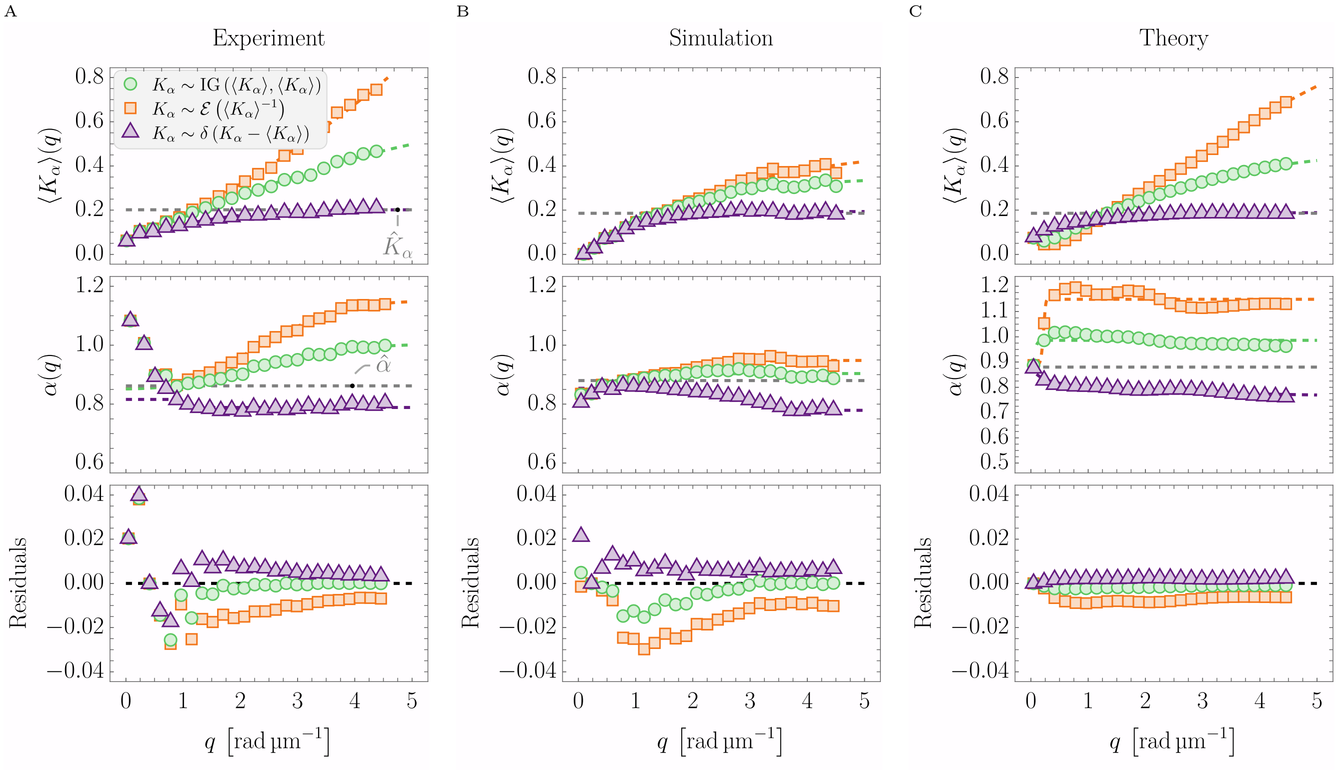


Figure S3: Comparison of the fitting parameters and residuals versus displacement mode $q$, obtained by fitting each closed-form ISF model to experimental data A, simulated data B, and numerical evaluation of (S11) C. For B and C we assumed inverse Gaussian distributed diffusivities and used the phenomenological form factor $F(q, R) \propto R^2/[1 + (qR)^2/2]$; $K_\alpha$ and $\alpha$ were chosen to match the experimental values. The residuals displayed in the bottom row were averaged over the lag time $\Delta t$ for each $q$. Across all panels, the inverse Gaussian model (green) has residuals closest to zero, while the exponential (orange) and delta (purple) models display larger, systematic deviations, most evident at lower $q$-values.

#### Microtubule segmentation and pore-size radius estimate

To obtain a radius estimate for the MT mesh size under stabilization, we segmented the MT network in each field of view (FOV) using custom `Mathematica` scripts and extracted a characteristic void length scale, see Figure S9. For each acquired FOV, we first averaged the images over all acquired frames and over the repeated $z$-stacks to obtain a single mean image, Figure S9A. Filamentous MT signal was enhanced with `RidgeFilter` and converted into a binary MT mask by adaptive thresholding. The binary mask was then skeletonized using `Thinning` and dilated with a disk structuring element to obtain a conservative representation of the MT network, which is shown as an overlay in Figure S9B.

Pores were defined as connected components of the complement of the MT mask, excluding components that touch the image boundary. For each pore, we computed the Euclidean distance transform of the pore mask and extracted the maximal distance within that pore. This maximum distance, denoted $\ell$, corresponds to the radius of the largest inscribed disk of the pore in 2D and serves as our pore-size radius estimate. Pooling $\ell$ over all pores yields the distribution $\mathcal{P}(\ell)$ shown in Figure S9C. We define $\hat{\ell}$ as the mode of the pooled distribution, and the FOV-resolved values of $\hat{\ell}$ are summarized in Figure S9D.

### Additional Analyses of Living HeLa Cells

#### Nuclear and Cytoplasmic Contributions

To assess how different cellular compartments contribute to the whole-cell DDM signal, we separately analyzed nuclear and cytoplasmic regions under control conditions. Because no fluorescent nuclear marker was available during the DDM acquisition, nuclei were manually delineated from the phase-contrast images. Cells for which the nuclear boundary could not be identified with sufficient contrast were excluded from this subcellular comparison. For each remaining cell, the cytoplasmic region was defined as the whole-cell mask minus the nuclear mask.

The resulting DDM analysis is shown in Figure S10. The fractional exponents are in agreement across the three masks, with $\langle \hat{\alpha}^{\mathrm{Whole}} \rangle = 0.813 \pm 0.007$, $\langle \hat{\alpha}^{\mathrm{Nuc}} \rangle = 0.807 \pm 0.009$, and $\langle \hat{\alpha}^{\mathrm{Cyt}} \rangle = 0.826 \pm 0.008$ (weighted average ± SEM). By contrast, the diffusivity is lower in the nucleus and higher in the cytoplasm than in the whole-cell analysis: $\langle \hat{K}_\alpha^{\mathrm{Whole}} \rangle = (0.973 \pm 0.030) \times 10^{-3}$, $\langle \hat{K}_\alpha^{\mathrm{Nuc}} \rangle = (0.483 \pm 0.018) \times 10^{-3}$, and $\langle \hat{K}_\alpha^{\mathrm{Cyt}} \rangle = (1.140 \pm 0.034) \times 10^{-3}$ (weighted average ± SEM). The whole-cell estimate therefore lies between the nuclear and cytoplasmic estimates, consistent with their combined contributions to the endogenous optical signal. The differences between the whole-cell and cytoplasm-only analyses remain comparatively small, and both indicate subdiffusive dynamics that are substantially slower than in egg extract.

Stronger compartment-specific differences may become apparent with imaging modalities that provide higher contrast and spatial resolution, such as DIC or phase-contrast microscopy with a higher-NA objective. Here, a comparatively low-NA air objective was deliberately used to establish the living-cell proof of principle.

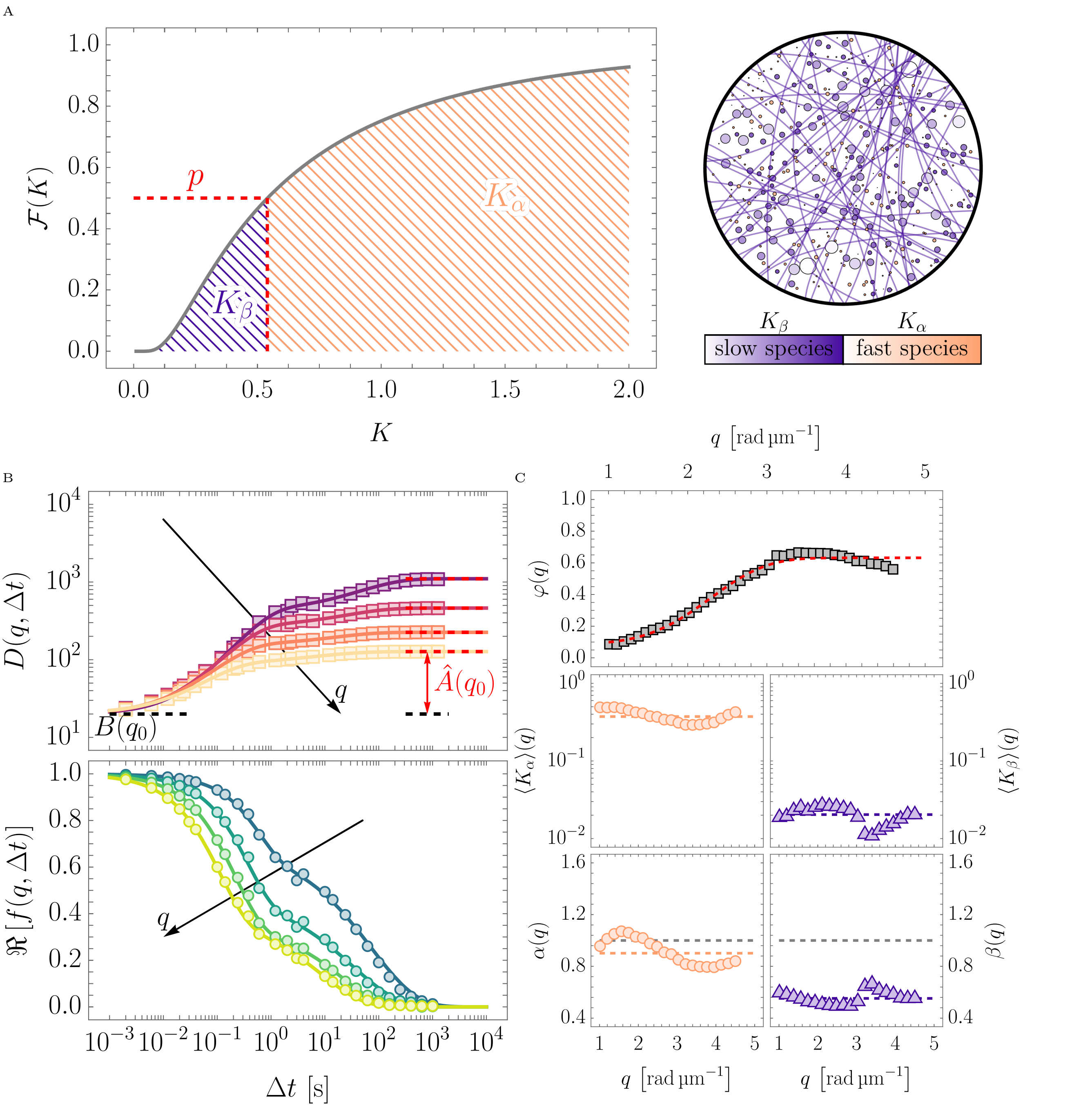


Figure S4: Overview and results of the two-state process simulation scheme described in the main text; A left: Cumulative distribution function of diffusivities $\mathcal{F}(K)$ of the inverse Gaussian distribution IG $(\langle K\rangle, \langle K\rangle)$ for $\langle K\rangle = 0.8\,\mu\text{m}^2/\text{s}$ (solid gray line) and the probability $p$ that defines the cut-off diffusivity $K^*$ (red dashed line) which is used to define $K \to K_\alpha$ for $K \geq K^*$ and $K \to K_\beta/10$ for $K < K^*$ with the respective fractional exponents $\alpha$ and $\beta$. A right: Schematic display of the simulated system, where the orange disks indicate the 'fast species' corresponding to the fractional Brownian motion process with diffusivity distribution $K \sim K_\alpha$ and fractional exponent $\alpha$, and the 'slow species' indicated by the purple disks and streaks corresponding to the fractional Brownian motion process with diffusivity distribution $K \sim K_\beta$ and fractional exponent $\beta$. B: DDM results of a simulation of the two-state process depicted in A for the case of $p = 0.2$ in analogy to Fig. 3 in the main text; top: DDM observable $D(q, \Delta t)$ and bottom: real part of the ISF $\Re[f(q, \Delta t)]$ (circles) and fits of the ISF using Eq. (9) of the main text. C: Fitting parameters of the data shown in B as a function of the displacement mode $q$; top: mixing parameter $\varphi$ where the red dashed line indicates the best fit to the data with a stretched sigmoidal function Eq. (10) in the main text, middle: dimensionally normalized diffusivities $K_\alpha$ (orange circles) and $K_\beta$ (purple up-triangles) where the dashed lines indicate the median values of the data, and bottom: fractional exponents $\alpha$ (orange circles) and $\beta$ (purple up-triangles) where the colorful dashed lines indicate the median of the data and the gray dashed lines indicate $\alpha = \beta = 1$.

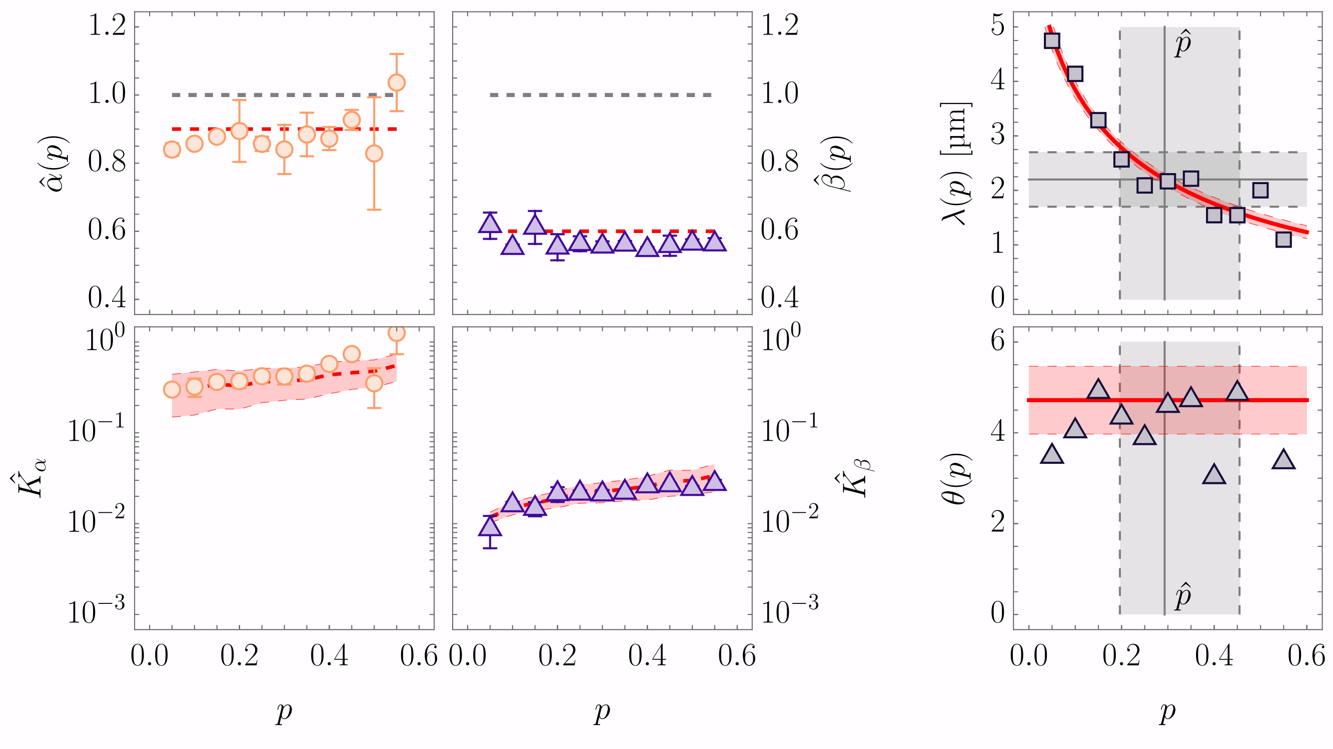


Figure S5: Results overview of the two-state simulation paradigm described in the text and shown in Figure S4; left: dimensionally normalized fractional diffusivity estimates $\hat{K}_\alpha$ and $\hat{K}_\beta$ (top row) and fractional exponent estimates $\hat{\alpha}$ and $\hat{\beta}$ (bottom row) as a function of the cut-off probability $p$, where the red dashed lines indicate the simulation input values (median with 68% CI) and the gray dashed lines for the fractional exponents indicate $\alpha = \beta = 1$. Right: Fitting results of $\varphi(q)$ with Eq. (10) in the main text over the cut-off probability $p$; characteristic length scale estimate $\hat{\lambda}$ (top) and scaling exponent $\hat{\theta}$ (bottom), where the red lines indicate best fit estimates ($\lambda(p) = \lambda_0 \exp[-(p/p_0)^{0.5}]$, and $\hat{\theta}(p) = \theta_0$) with 68% CIs. The gray dashed lines indicate the experimental $\hat{\lambda}$ of HSS+Vanadate+TAX (see main text; median ± 68% CIs) and the inferred $\hat{p}$ using the fit $\lambda(p)$.

### Apparent Cell Motility During DDM Acquisition

To assess whether whole-cell motion contributes appreciably to the measured ISFs, the positions of segmented cells were evaluated throughout the acquisition. For this quantification, cells were segmented every 100 frames at a frame rate of 5 Hz. At each time point, centroid displacements were summarized by the median across cells within each field of view and then averaged over three fields of view.

As shown in Figure S11, the mean displacement remains well below 1 µm over the approximately 1000 s measurement window in both conditions. With a pixel size of approximately 0.10 µm px$^{-1}$. Whole-cell migration is therefore a negligible contribution to the reconstructed dynamics over the analyzed lag-time and spatial frequency range.

### Further Materials

Here we present additional figures that corroborate the findings presented in the main text.

Figure S12 provides fluorescence images of the cell conditions studied in Fig. 4 in the main text. Control cells show an intact MT network, while nocodazole treatment reduces and disrupts the network.

Figure S13 depicts the noise characteristics of the camera used in this study in terms of the background DDM observable $\langle D_0(\langle I_0\rangle, q_0, \Delta t)\rangle_{\Delta t}$ for different mean background intensities $\langle I_0\rangle$, averaged over lag time at a fixed displacement mode $q_0$. The resulting curve was used to select an intensity range that optimizes dynamic range while avoiding under- and overexposure, as described in the main text.

Figure S14 shows exemplarily the $q$-dependence of the relative heterogeneity $\kappa(q)$ for high-speed egg extract.

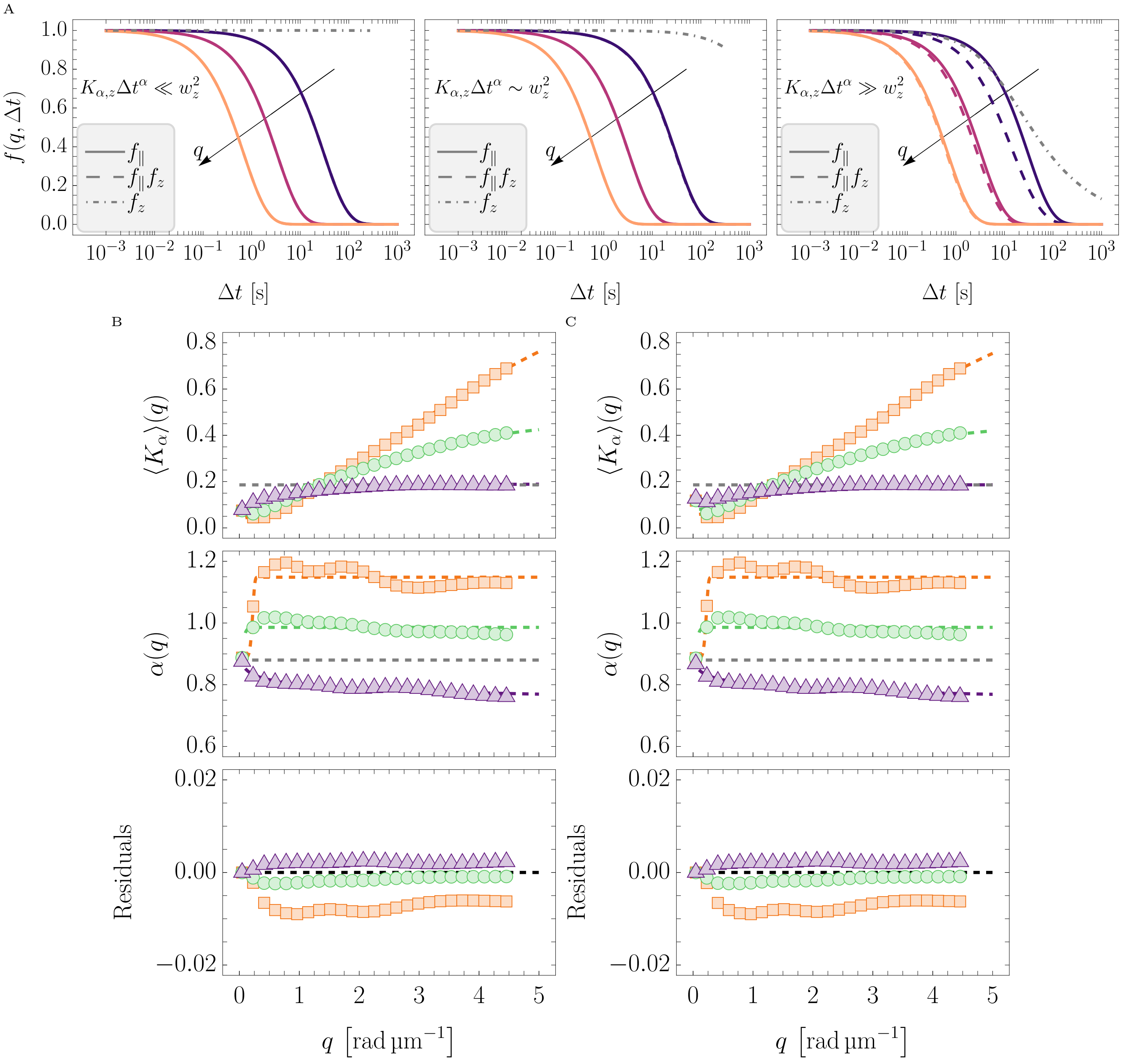


Figure S6: Effect of axial decorrelation on the ISF and on the fitted DDM parameters. A: In-plane ISF $f_\parallel(q, \Delta t)$ (solid lines), axial decorrelation factor $f_z(\Delta t)$ from Equation (S29) (dash-dotted lines), and product $f_\parallel(q, \Delta t) f_z(\Delta t)$ (dashed lines) for selected displacement modes $q$. The three cases illustrate regimes where $K_{\alpha,z}(\Delta t)^\alpha / w_z^2$ remains small (left), becomes comparable to one (middle), or exceeds one (right). Axial diffusion affects the observed ISF only when this ratio approaches or exceeds unity, with the strongest effect at low $q$, where in-plane relaxation is slow. B and C: Fitting parameters and residuals versus $q$, obtained by fitting each closed-form ISF model (inverse Gaussian: green circles; exponential: orange squares; delta: purple triangles) to the numerical evaluation of Equation (S11) without B and with C the axial decorrelation term. We assumed inverse Gaussian distributed diffusivities and used the phenomenological form factor $F(q, R) \propto R^2/[1 + (qR)^2/2]$; $K_\alpha$ and $\alpha$ were chosen to match the experimental values of HSS (gray dashed lines). The sectioning width $w_z \simeq \lambda^2/\Delta\lambda \approx 20\,\mu\mathrm{m}$ was estimated from the LED spectral coherence, and $K_{\alpha,z} = K_\alpha$ was used as a conservative upper estimate. Fitting residuals in the bottom row were averaged over $\Delta t$ for each $q$.

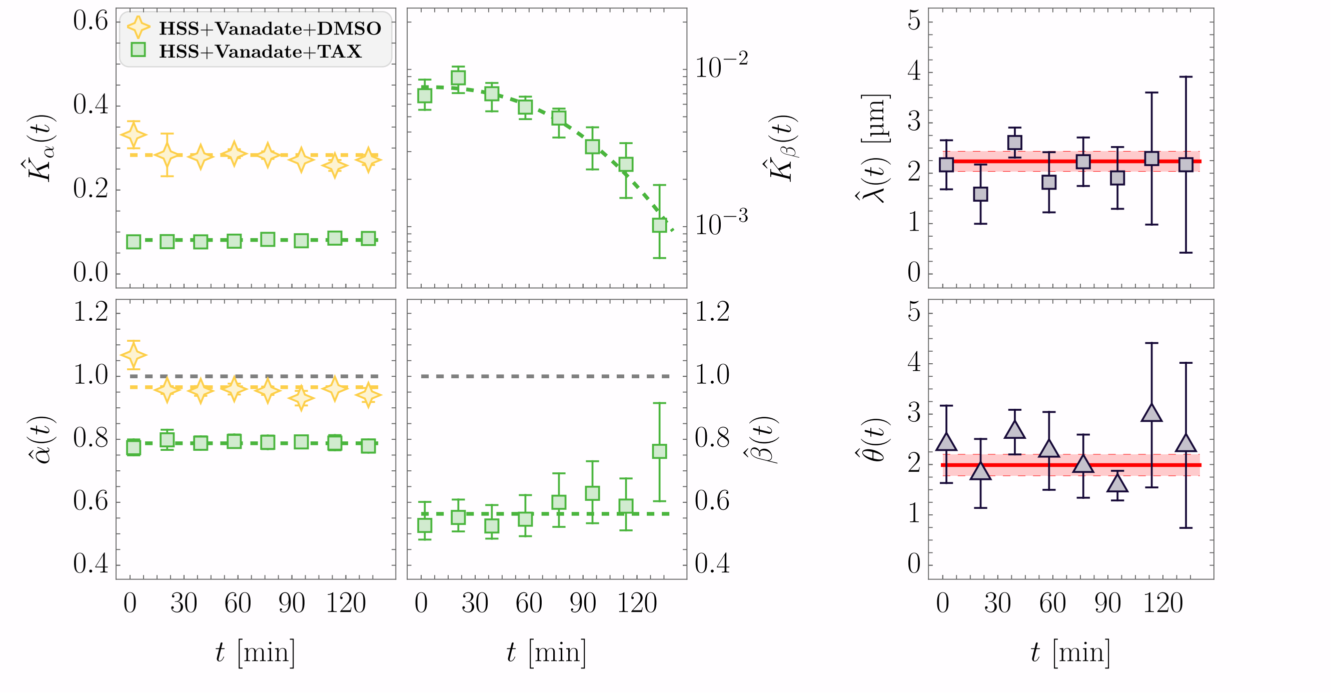


Figure S7: Results of the DDM analysis of time-series measurements of HSS with Vanadate (0.5 mM) and Taxol (0.25 µM): Left: Dimensionally normalized fractional diffusivity estimates $\hat{K}_\alpha$ and $\hat{K}_\beta$ (top row), and fractional exponent estimates $\hat{\alpha}$ and $\hat{\beta}$ (bottom row) for HSS+Vanadate+Taxol (green squares) and control HSS+Vanadate+DMSO (yellow stars) as a function of measurement time $t$. Colorful dashed lines are best fit estimates as in Fig 3 in the main text; gray dashed lines for fractional exponents indicate $\alpha = \beta = 1$. Right: Fitting results for $\varphi(q)$ with Eq. (10) in the main text as a function of measurement time $t$: characteristic length scale estimate $\hat{\lambda}$ (top) and scaling exponent estimate $\hat{\theta}$ (bottom), with red lines as constant best fit estimates and 68% CIs. Each data point shows mean ± pooled SEM of three independent measurements (see *Materials and Methods*).

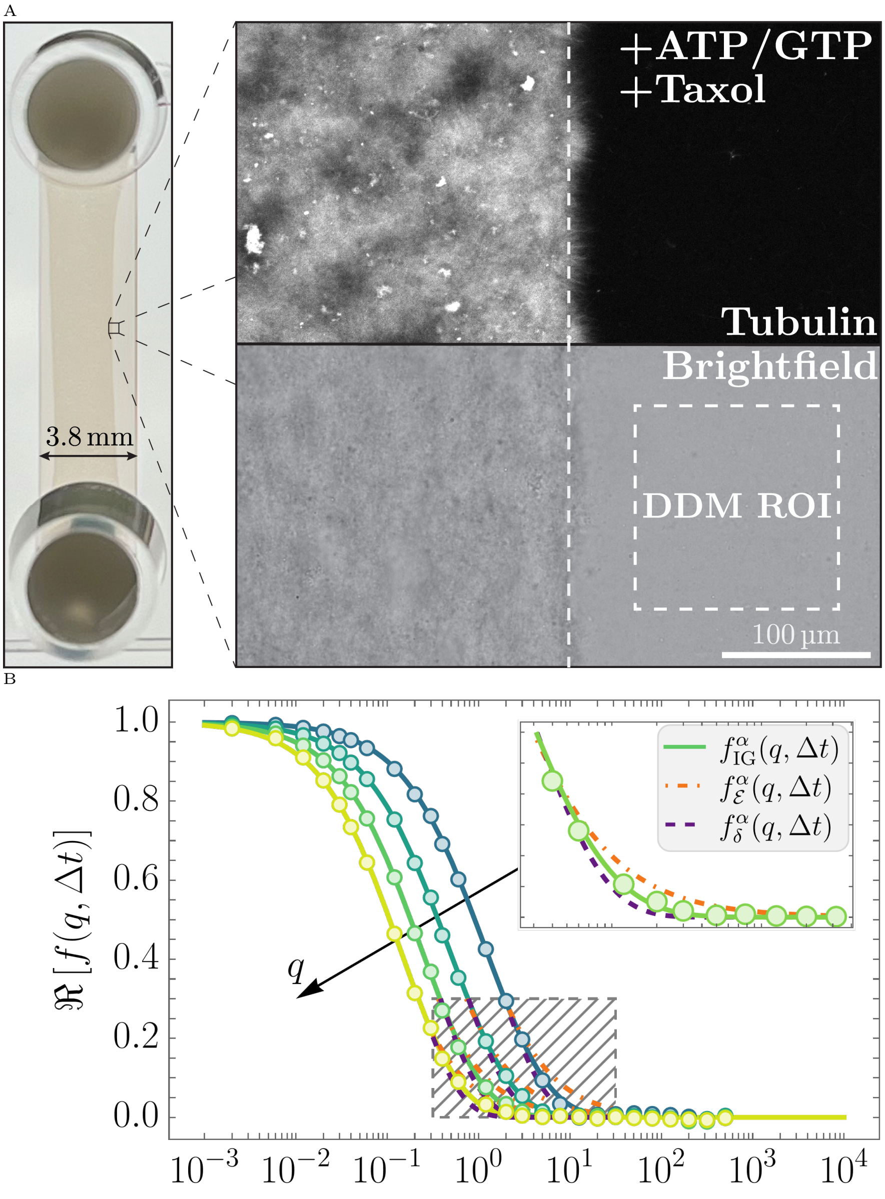


Figure S8: Results of the DDM analysis of HSS with the addition of 20x Energy Mix and Taxol (5 µM): A: Top-view image of the mounting channel approximately 2.5 h after preparation with a clearly visible MT contraction band (left) and a corresponding confocal fluorescent (HyLyte 488-labeled tubulin) and bright-field microscopy images of the region indicated by the black square (right). B: Real part of the ISF $\Re\,[f(q,\Delta t)]$ as a function of the lag time $\Delta t$ and displacement modes $q$, obtained with DDM of the region of interest indicated by the dashed white square in A (right).

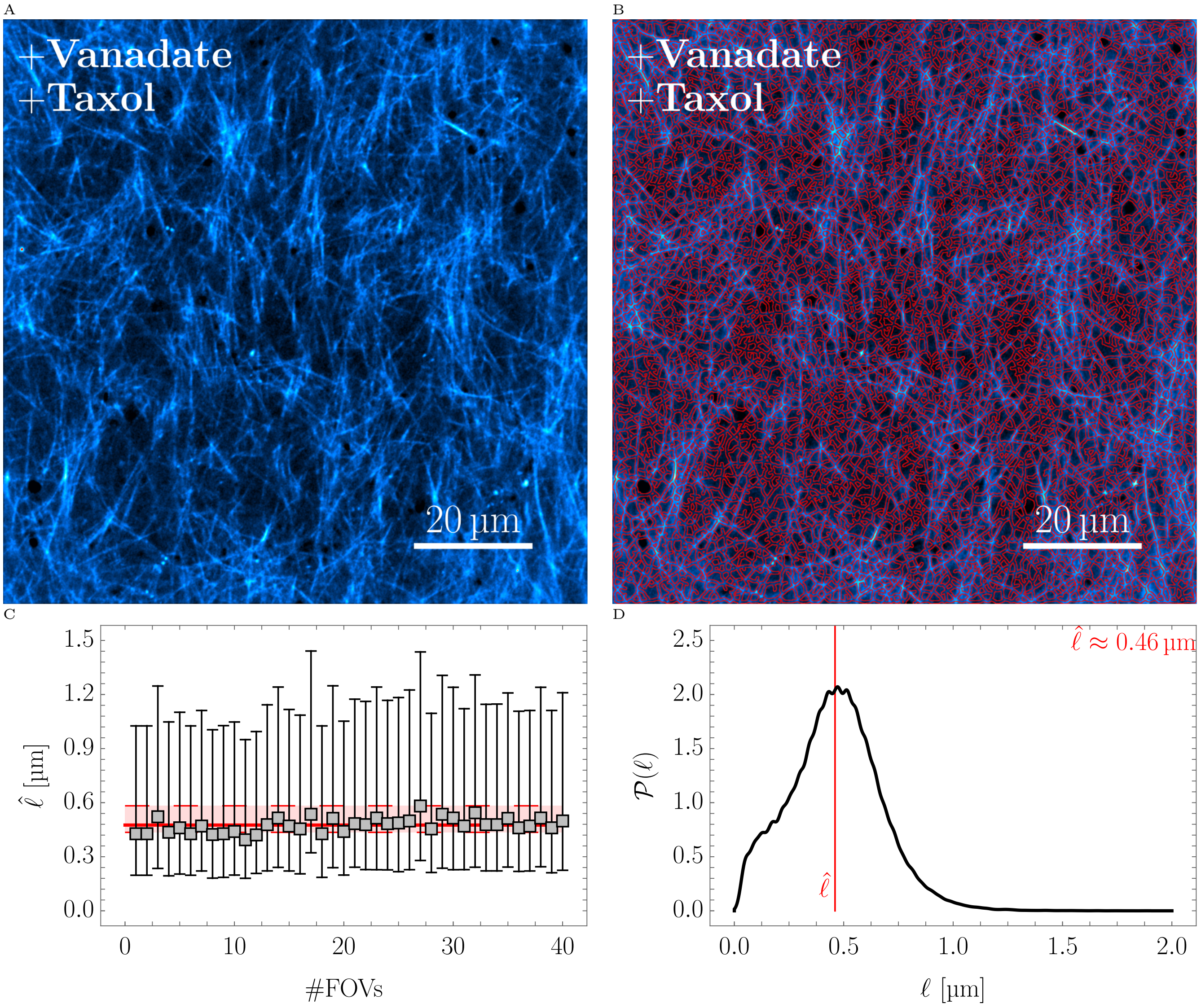


Figure S9: MT segmentation and pore-size radius estimate in Taxol-stabilized extracts. A: Representative average MT image for one FOV. B: Segmentation overlay showing the MT network mask obtained from `RidgeFilter`-based enhancement, skeletonization, and dilation with a disk structuring element. C: Pore-size radius estimate $\hat{\ell}$ per FOV, where each marker denotes the median $\ell$ within one FOV and the error bars indicate the central 68% interval; the red line and shaded band show the pooled estimate and its uncertainty. D: Pooled distribution $\mathcal{P}(\ell)$ from all pores across all FOVs, with the red line indicating the mode $\hat{\ell}$.

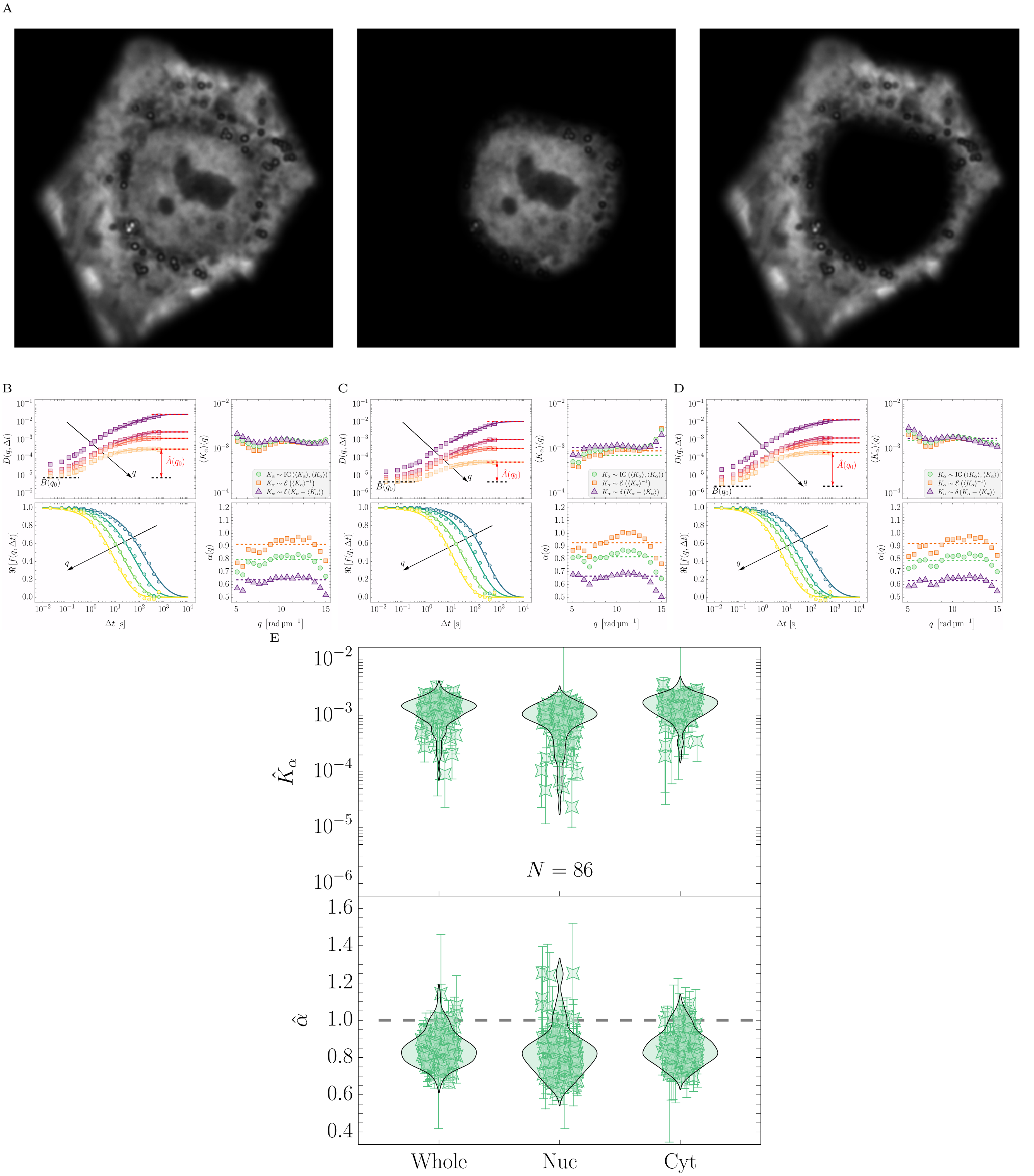


Figure S10: Compartment-resolved DDM analysis of living HeLa cells under control conditions. A: Representative phase-contrast image with masks for the whole cell (left), nucleus (middle), and cytoplasm (right). B–D: DDM results, in analogy to Fig. 4 in the main text, for the whole cell, nucleus, and cytoplasm, respectively. E: Summary of the best-estimate parameters $\hat{K}_\alpha$ and $\hat{\alpha}$ across control cells (HeLa+DMSO) for whole-cell, nuclear, and cytoplasmic masks. Points show individual cells as median ± 68% CI, and violin outlines indicate the corresponding distributions.

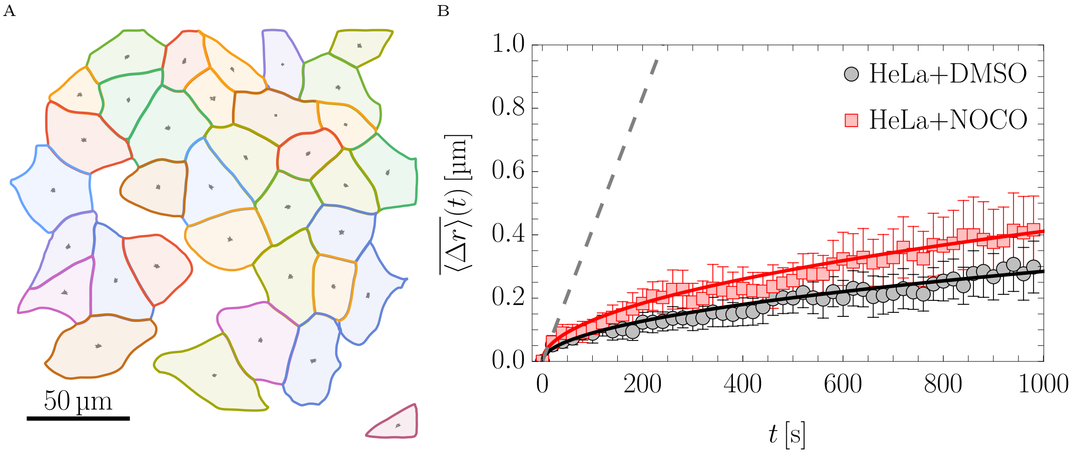


Figure S11: Apparent cell motility over time. A: Representative segmentation mask with centroid tracks over 1000 s (gray). B: Corresponding mean displacement $\overline{\langle \Delta r \rangle}(t)$ of HeLa cells over time under control conditions (DMSO, black circles) and after Nocodazole treatment (red squares). Symbols show the mean over three fields of view at each time point, and error bars indicate the corresponding standard deviation. For each field of view, the median displacement and median deviation were evaluated over all segmented cells. Solid lines indicate $\overline{\langle \Delta r \rangle}(t) \propto \sqrt{t}$, while the gray dashed line shows $\overline{\langle \Delta r \rangle}(t) = 15\,\mu\mathrm{m}\,\mathrm{h}^{-1} t$, corresponding to the migration velocity reported for sparse HeLa cells in [18], for visual guidance.

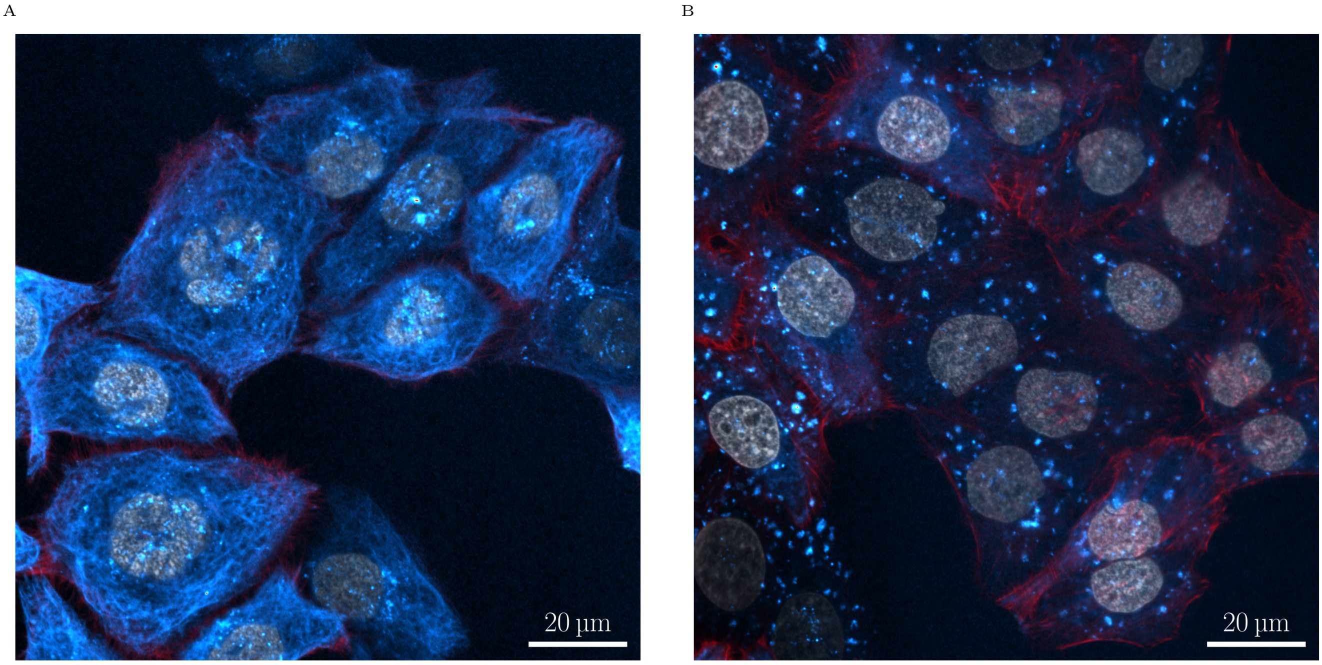


Figure S12: Fluorescence images of HeLa cells under different conditions. A: DMSO control cells with F-actin (red), $\alpha$-tubulin (blue), and DAPI (gray). B: Nocodazole-treated cells (HeLa+NOCO) with the same markers, showing reduced and disorganized MT signal compared to control.

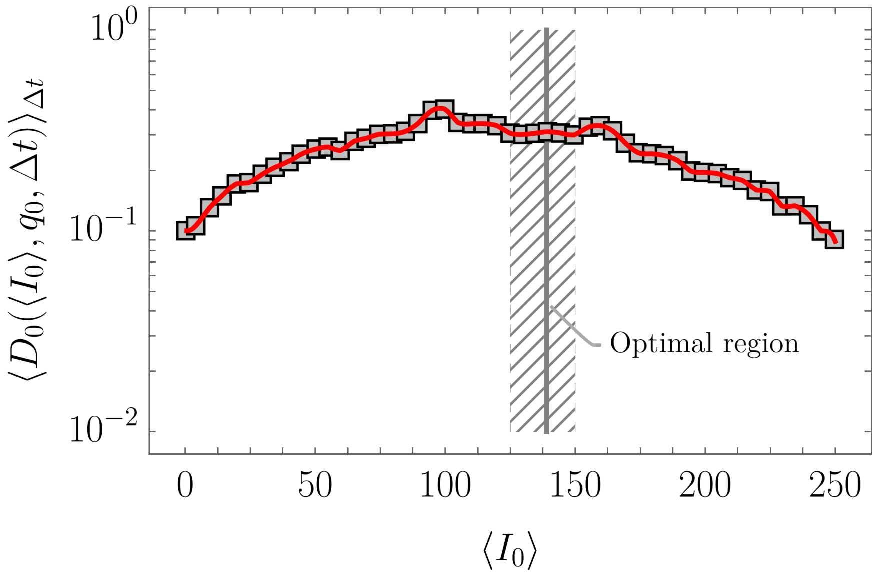


Figure S13: Noise characteristics of the Mikrotron EoSens CMOS sensor used in this study. The plot shows the average background DDM map $\langle D_0(\langle I_0 \rangle, q_0, \Delta t) \rangle_{\Delta t}$ for a specific displacement mode $q_0 \approx 4.13\,\mathrm{rad}\,\mu\mathrm{m}^{-1}$ at various mean intensities $\langle I_0 \rangle$ (black squares) alongside the corresponding interpolation function (red curve). The gray hatched area delineates the optimal mean intensity range for measurements ($125 \leq \langle I \rangle \leq 150$), and the solid gray line indicates the optimized mean intensity used in this study.

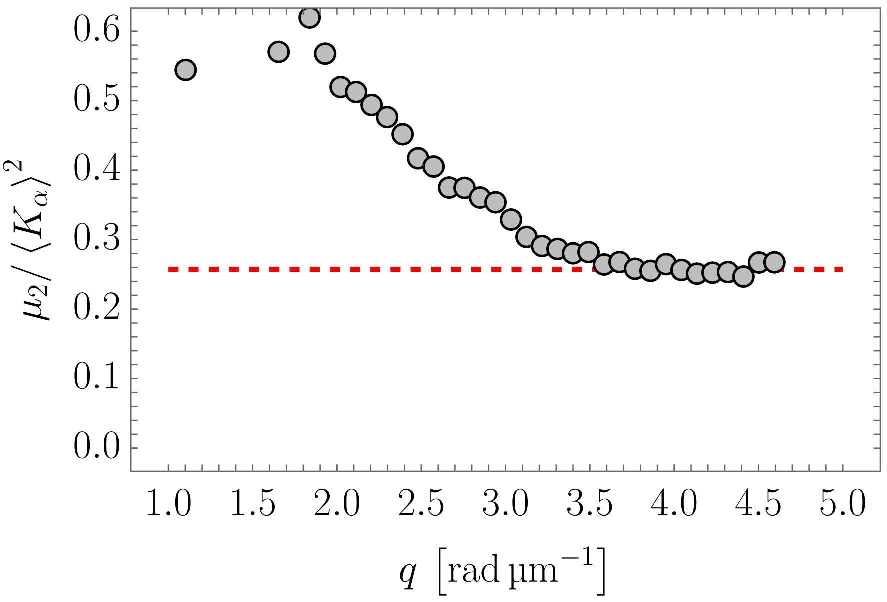


Figure S14: Relative heterogeneity parameter $\kappa \equiv \mu_2/\langle K_\alpha \rangle^2$ (see Equation (S24)) in dependence of the displacement mode $q$ for high-speed egg extract. The red dashed line indicates the median value for $3.5\,\mathrm{rad\,\mu m^{-1}} \leq q \leq \pi/\epsilon$ (see *Materials and Methods* for more information).
